# Genome-wide association across *Saccharomyces cerevisiae* strains reveals substantial variation in underlying gene requirements for toxin tolerance

**DOI:** 10.1101/241257

**Authors:** Maria Sardi, Vaishnavi Paithane, Michael Place, De Elegant Robinson, James Hose, Dana J. Wohlbach, Audrey P. Gasch

## Abstract

Cellulosic plant biomass is a promising sustainable resource for generating alternative biofuels and biochemicals with microbial factories. But a remaining bottleneck is engineering microbes that are tolerant of toxins generated during biomass processing, because mechanisms of toxin defense are only beginning to emerge. Here, we exploited natural diversity in 165 *Saccharomyces cerevisiae* strains isolated from diverse geographical and ecological niches, to identify mechanisms of hydrolysate-toxin tolerance. We performed genome-wide association (GWA) analysis to identify genetic variants underlying toxin tolerance, and gene knockouts and allele-swap experiments to validate the involvement of implicated genes. In the process of this work, we uncovered a surprising difference in genetic architecture depending on strain background: in all but one case, knockout of implicated genes had a significant effect on toxin tolerance in one strain, but no significant effect in another strain. In fact, whether or not the gene was involved in tolerance in each strain background had a bigger contribution to strain-specific variation than allelic differences. Our results suggest a major difference in the underlying network of causal genes in different strains, suggesting that mechanisms of hydrolysate tolerance are very dependent on the genetic background. These results could have significant implications for interpreting GWA results and raise important considerations for engineering strategies for industrial strain improvement.

**Author summary:** Understanding the genetic architecture of complex traits is important for elucidating the genotype-phenotype relationship. Many studies have sought genetic variants that underlie phenotypic variation across individuals, both to implicate causal variants and to inform on architecture. Here we used genome-wide association analysis to identify genes and processes involved in tolerance of toxins found in plant-biomass hydrolysate, an important substrate for sustainable biofuel production. We found substantial variation in whether or not individual genes were important for tolerance across genetic backgrounds. Whether or not a gene was important in a given strain background explained more variation than the alleleic differences in the gene. These results suggest substantial variation in gene contributions, and perhaps underlying mechanisms, of toxin tolerance.

## Introduction

The increased interest in renewable energy has focused attention on non-food plant biomass for the production of biofuels and biochemicals [1]. Lignocellulosic plant material contains significant amounts of sugars that can be extracted through a variety of chemical pretreatments and used for microbial production of alcohols and other important molecules [2–5]. However, there are major challenges to making biofuel production from plant biomass economically viable [6]. One significant hurdle with regards to microbial fermentation is the presence of toxic compounds in the processed plant material, or hydrolysate, including weak acids, furans and phenolics released or generated by the pretreatment process [7–10]. The concentrations and composition of these inhibitors vary for different pretreatment methods and depend on the plant feedstocks [7, 9, 11]. These toxins decrease cell productivity by generating reactive oxygen species, damaging DNA, proteins, cell membranes [12–14], and inhibiting important physiological processes, including enzymes required for fermentation [15], *de novo* nucleotide biosynthesis [16], and translation [17]. Despite knowledge of these targets, much remains to be learned about how the complete suite of hydrolysate toxins (HTs) acts synergistically to inhibit cells. Furthermore, how the effects of HTs are compounded by other industrial stresses such as high osmolarity, thermal stress, and end-product toxicity remains murky.

Engineering strains with improved tolerance to industrial stresses including those in the plant hydrolysate is of the utmost importance for making biofuels competitive with fuels already in the market [6]. A goal in industrial strain engineering is to improve lignocellulosic stress tolerance, often through directed engineering. Many approaches have been utilized to identify genes and processes correlated with increased stress tolerance, including transcriptomic profiling of cells responding to industrial stresses [18–21], genetic mapping in pairs of strains with divergent phenotypes [22–25], and directed evolution to compare strains selected for stress tolerance with starting strains [26–29]. However, in many cases the genes identified from such studies do not have the intended effect when engineered into different genetic backgrounds [30–33]. One reason is that there are likely to be substantial epistatic interactions between the genes identified in one strain and the genetic background from which it was identified [34]. A better understanding of how tolerance mechanisms vary across genetic backgrounds is an important consideration in industrial engineering.

Exploring variation in HT tolerance across strain background could also reveal additional defense mechanisms. The majority of functional studies in *Saccharomyces cerevisiae* are carried out in a small number of laboratory strains that do not represent the rich diversity found in this species [35, 36]. The exploration of natural diversity in *S*. *cerevisiae* has revealed a wide range of genotypic and phenotypic variability within the species [36–40]. In some cases, trait variation is correlated with genetic lineage [36, 41–43], indicating a strong influence of population history. At least 6 defined lineages have been identified in the species, including strains from Malaysia, West Africa, North America, Europe/vineyards, and Asia [41] as well as recently identified populations from China [38, 44]. In addition to genetic variation, phenotypic variation has cataloged natural differences across strains, in transcript abundance [37, 45, 46], protein abundance [47–49], metabolism [50–52], and growth in various environments [32, 36, 37, 42, 52–54]. Thus, *S*. *cerevisiae* as a species presents a rich resource for dissecting how genetic variation contributes to phenotypic differences. In several cases this perspective has benefited industry in producing novel strains by combining genetic backgrounds or mapping the genetic basis for trait differences [25, 55–59].

We used genome-wide association (GWA) in *S*. *cerevisiae* strains responding to synthetic hydrolysate (SynH), both to identify new genes and processes important for HT tolerance and to explore the extent to which genetic background influences mechanism. We tested 20 genes associated with HT tolerance and swapped alleles across strains to validate several allele-specific effects. However, in the process of allele exchange we discovered striking differences in gene contributions to the phenotype: out of 14 gene knockouts tested in two strains with opposing phenotypes, 8 (57%) had a statistically significant effect on HT tolerance in one of the backgrounds but little to no significant effect in the other background. In most of these cases, the specific allele had little observable contribution to the phenotype. Thus, although GWA successfully implicated new genes and processes involved in HT tolerance, the causal variation in the tested strains is not at the level of the allele but rather whether or not the gene’s function is important for the phenotype in that background. This raises important implications for considering natural variation in functional networks to explain phenotypic variation.

## Results

### Genetic variation across 165 *S*. *cerevisiae* strains

We obtained 165 *Saccharomyces cerevisiae* strains, representing a range of geographical and ecological niches, that have high quality whole genome sequencing reads (coverage ~30X), coming from published sequencing projects across the yeast community [39, 42, 52, 60] (S1 Table). We identified 486,302 high quality SNPs (see Methods). 68% of them had a minor allele frequency less than 5%. Nucleotide variation compared to the well-studied S288c-derived reference strain varied from as low as 0.08% for the closely related W303 lab strain and as high as 0.72% for the bakery strain YS4 (S1 Table). The majority of strains were largely homozygous (in some cases due to strain manipulation by sequencing projects); however, we identified 21 strains with >20% heterozygous sites. Most of these were from natural environments (11 strains) but they also included clinical samples (5 strains), baking strains (3 strains), a sugar cane fermenter (1) and a laboratory strain (FL100, which was scored as 98% heterozygous and may have mated with another strain in its recent history (S1 Table)).

Sixty-three percent of the variants were present in coding regions (S2 Table), which is lower than random expectation (since 75% of the yeast genome is coding) and consistent with purifying selection acting on most gene sequences. Indeed, coding variants predicted to have high impact, such as SNPs that introduce a stop codon, eliminate the start codon, or introduce a defect in the splicing region, were very rare (0.004% of genic SNPs) - a third of these were in dubious ORFs (22%) or genes of unknown function (8%) [61] that are likely nonfunctional and under relaxed constraint. However, 54 genes with debilitating polymorphisms are reportedly essential in the S288 background; nearly half of these polymorphisms are present in at least 3 strains and in some cases are lineage specific (S3 Table). Tolerance of these polymorphisms could arise through duplication of a functional gene copy [62], but could also arise due to evolved epistatic effects as has been previously reported [63], highlighting the complexity behind genetic networks and the role of genetic variation in determining their regulation.

Principle component analysis of the genomic data recapitulated the known lineages represented in the collection, including the European/wine, Asian/sake, North American (NA), Malaysian, West African (WA), and mosaic groups [36, 41, 42, 64] (S1 Table). Our analysis split the West African population into three subgroups not previously defined (Fig 1A). Construction of a simple neighbor-joining (NJ) tree broadly confirmed the population groups present in the 165-strain collection (Fig 1 B).

**Fig 1.**
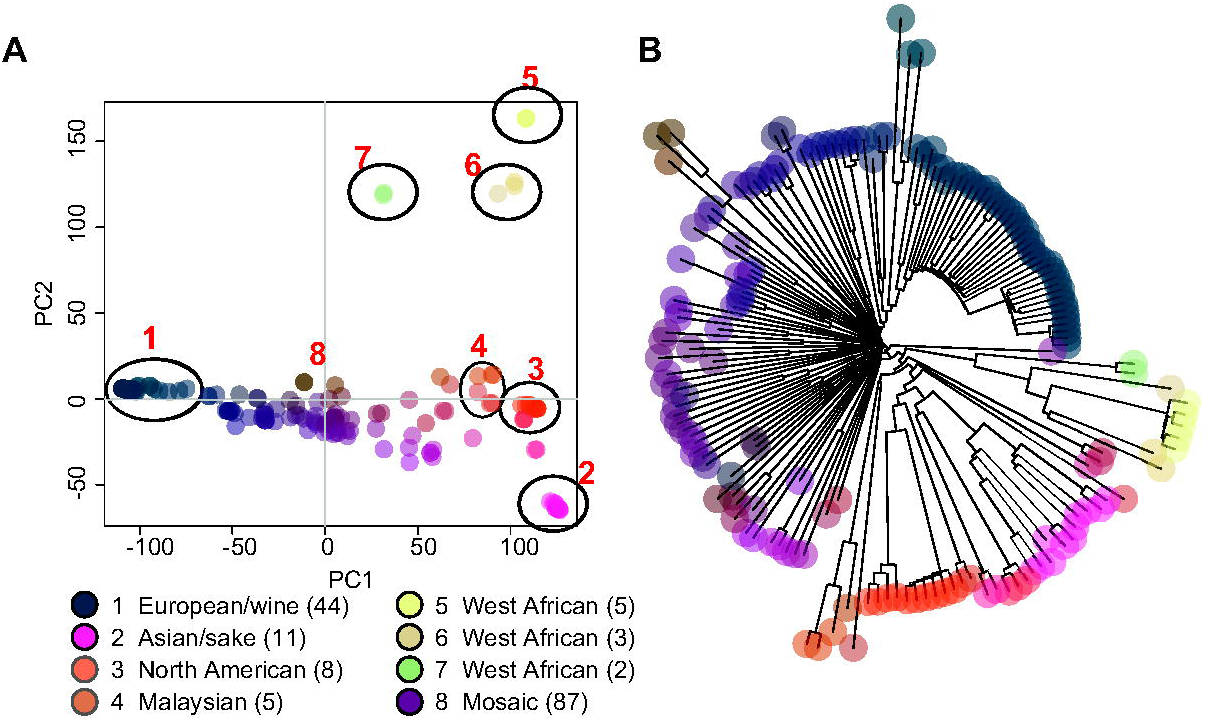
Genetic diversity found in the 165-strain collection. The entire collection of high quality SNPs (486,302) was used as input for principal component analysis (PCA) (A) and to generate a neighbor-joining (NJ) tree (B). Populations assigned in circles were defined manually using published population structure data. Strains are color-coded according to genetic similarity, with matching colors between the PCA and NJ tree (generated by Adegenet [110]). The population and/or niche is represented by the key, with the number of strains in each group indicated in parentheses.

### Phenotypic variation in SynH tolerance is partly correlated with ancestral group

We scored variation in lignocellulosic hydrolysate tolerance in several ways. Strains that are sensitive to hydrolysate grow slower and consume less sugars over time [65], thus we measured final cell density and percent of glucose consumed after 24 hours to represent SynH tolerance. Growth and glucose consumption were significantly correlated (R^2^ = 0.79), although there was some disagreement for particular strains (including flocculant strains) (S1 Table). We also determined tolerance to HTs specifically, to distinguish stress inflected by HTs from effects of the base medium that has unusual nutrient composition and high osmolarity due to sugar concentration. To do this, we calculated the relative percent-glucose consumed and final OD_600_ in media with (SynH) and without HT toxins (SynH–HTs, see Methods) (S1 Table). Tolerance to SynH base medium without toxins (SynH–HT) and SynH with the toxins was only partly correlated (R^2^ = 0.48) (S1 Fig), suggesting that there are separable mechanisms of growing in base medium and surviving the toxins.

There is wide variation in tolerance to lignocellulosic hydrolysate that partly correlates with populations (Fig 2, S1 Fig). North American and Malaysian strains displayed the highest tolerance to SynH. As expected, phenotypic variation within each population was related to genetic variation, *e.g*. West African strains in Population 5 showed low genetic and phenotypic variation while mosaic strains with genetic admixture showed the widest range of phenotypes.

**Fig 2.**
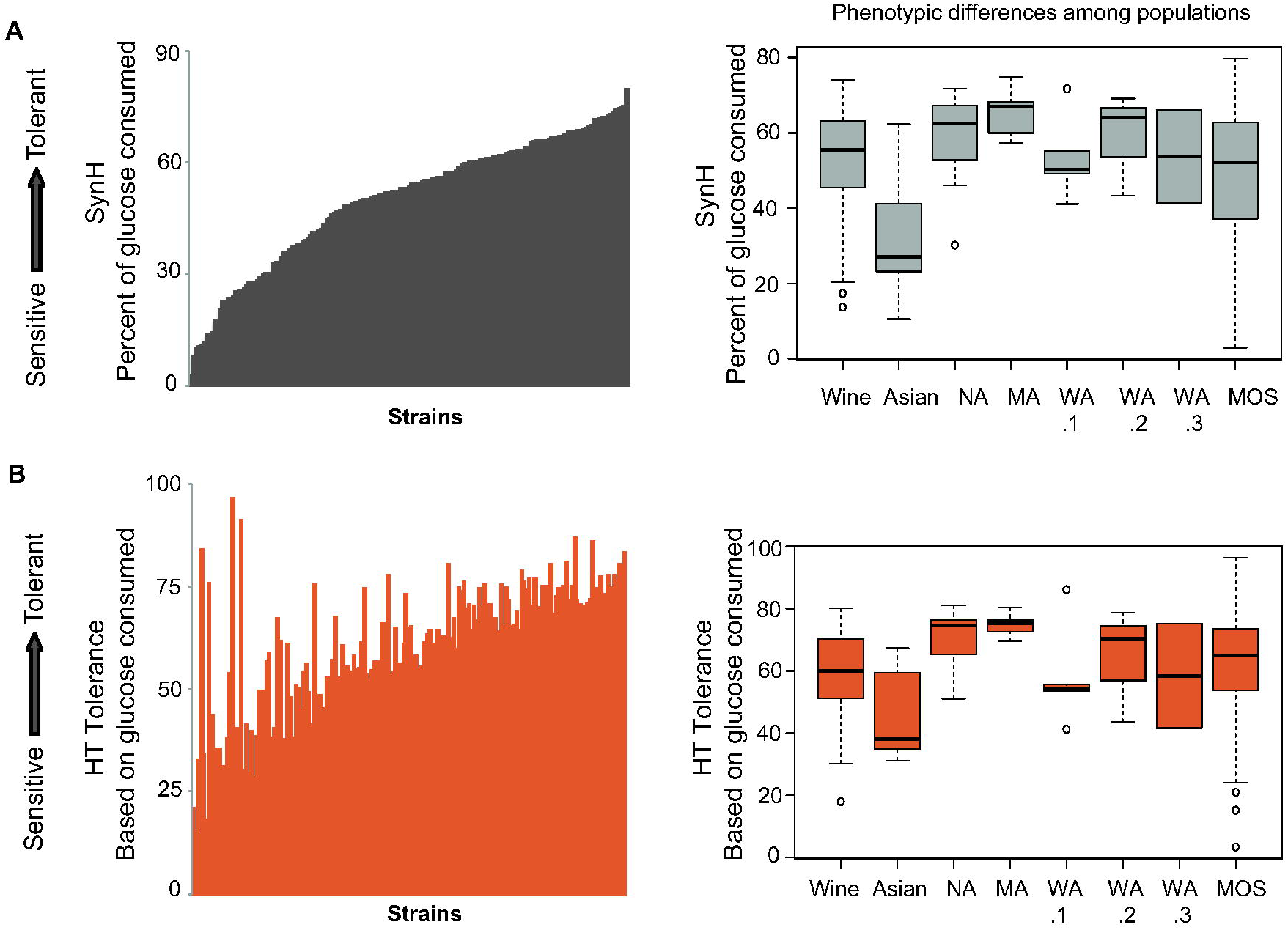
Strain-specific difference for SynH and HT tolerance. Tolerance to lignocellulosic hydrolysate across strains (left) and across each population (right) measured as glucose consumption in SynH (A) and HT tolerance based on glucose consumption were calculated as described in Methods for 165 strains. Individual strains were ordered based on the quantitative scores in (A). Population distributions shown in the boxplots are indicated for named populations from Fig 1.

### Genome-wide association analysis reveals genes associated with SynH tolerance

We used GWA to map the genetic basis for the differences in SynH tolerance, for each of the four phenotypes introduced above. The population signatures in *S*. *cerevisiae* are problematic for GWA, since the strong correlations between phenotype and ancestry obscure the identification of causal polymorphisms [66, 67]. To overcome this, we incorporated a large number of mosaic strains in the analysis and used a mixed-linear model to account for strain relationships, as implemented in the program GAPIT [68] (see Methods). We used as input SNPs that were present in at least 3 strains, eliminating 42% of SNPs in the dataset (see Methods). Of the remaining SNPs, 45% have a minor allele frequency of less than 5%; only those with an allele frequency >2% were used for GWA. GWA identified loci whose variation correlated with phenotypic variation. None of the GWA-implicated loci passed the stringent Bonferroni p-value correction based on the number of effective tests (see Methods), which is not uncommon for GWA at this scale [42, 69, 70]. We therefore used a somewhat arbitrary p-value cutoff of 1e-04 and performed additional filtering to minimize false positive associations (see Methods).

The combined analysis yielded 76 SNPs that met our p-value threshold (S4 Table, S2 Fig). Thirty-eight of these SNPs, linked to 33 genes, passed additional filtering (See Methods, Table 1). Of these, 17 SNPs are associated with growth in SynH, while 23 SNPs are associated with tolerance to HTs specifically (Table 1). Eight of the SNPs are intergenic and 20 are located within genes, with 13 of those predicted to change the coding sequence. Although we would expect that SNPs linked to HT tolerance should be identified in both sets of analyses, only 2 SNPs were significantly associated with both SynH and HT tolerance. This almost certainly highlights limited statistical power with the small set of strains used here. For most SNPs, the allele associated with tolerance was more frequent in our strain collection (Fig 3A), but for some it was the allele associated with sensitivity that was nearly fixed. We carried out additional GWA filtering to ensure that results were not driven by population structure (see Methods), since we note that many of sensitive alleles were prominent in the Asian population (S3 Fig). As expected for a largely additive trait, there was a significant linear correlation between the number of deleterious alleles a strain harbored and its tolerance to hydrolysate (R^2^ = 0.48, p = 2.2e-16, Fig 3B).

**Table 1.** SNPs associated with SynH tolerance.

| Phen. | Locus<br>Chr: Pos | Type | Gene/Region | Function | p Value |
| --- | --- | --- | --- | --- | --- |
| 4 | 12:1033361 | syn | HMG2 | HMG coA reductase | 3.20E-06 |
| 4 | 11:649062 | mis | FLO10 | flocculation protein | 9.30E-06 |
| 1 & 2 | 16:230703 | mis | MEX67 | mRNA export | 1.10E-05 |
| 3 | 16:897004 | splice site | AOS1 | SUMO E1 | 1.30E-05 |
| 1 | 04:122163 | int | UFD2 / | Ubiquitin assembly | 1.40E-05 |
| 1 | 04:122163 | int | RBS1 | RNA Pol II assembly | 1.40E-05 |
| 4 | 12:1034877 | mis | HMG2 | HMG coA reductase | 1.50E-05 |
| 3 | 15:993416 | syn | MNE1 | mitochondrial matrix p. | 1.70E-05 |
| 3 | 15:993550 | syn | MNE1 | mitochondrial matrix p. | 1.70E-05 |
| 3 | 15:993749 | syn | MNE1 | mitochondrial matrix p. | 1.70E-05 |
| 3 | 15:993770 | syn | MNE1 | mitochondrial matrix p. | 1.70E-05 |
| 4 | 15:983849 | syn | REV1* | DNA damage repair | 2.60E-05 |
| 1 | 11:152140 | int | KDX1* / | MAP kinase | 3.20E-05 |
| 1 | 11:152140 | int | ELF1 | transcription elongation | 3.20E-05 |
| 3 | 15:840176 | syn | RIM20 | transcription reglugator | 3.60E-05 |
| 2 | 13:685585 | syn | ERG12 | ergosterol synthesis | 3.60E-05 |
| 3 | 12:492470 | mis | MAS1 | mitochondrial protein import | 4.40E-05 |
| 1 & 4 | 12:1037554 | mis | LEU3 | transcription factor | 4.60E-05 |
| 4 | 02:161990 | mis | SHE1 | spindle protein | 4.80E-05 |
| 4 | 15:984317 | syn | REV1* | DNA damage repair | 4.90E-05 |
| 2 & 4 | 07:20180 | int | ADH4* / | alcohol dehydrogenase | 5.20E-05 |
| 2 & 4 | 07:20180 | int | ZRT1 | zinc transport | 5.20E-05 |
| 3 | 16:897179 | int | AOS1 / | SUMO E1 | 5.30E-05 |
| 3 | 16:897179 | int | SEC1 | secretion | 5.30E-05 |
| 1 & 2 | 09:333735 | syn | TIR3* | cell wall protein | 6.20E-05 |
| 4 | 14:341663 | syn | NSG2 | sterol biosynthesis | 6.30E-05 |
| 3 | 01:203973 | mis | FLO1* | flocculation protein | 6.70E-05 |
| 1 & 2 | 12:691485 | int | PIG1* / | regulates Glc7 phosphatase | 7.00E-05 |
| 1 & 2 | 12:691485 | int | MCM5 | DNA replication | 7.00E-05 |
| 3 | 13:599730 | syn | ALD3* | aldehyde dehydrogenase | 7.80E-05 |
| 2 | 16:406798 | intron | RPL21B* | ribosomal protein | 8.40E-05 |
| 4 | 13:44254 | syn | DAT1* | DNA binding protein | 8.50E-05 |
| 1 | 11:496123 | syn | SAP190 | phosphatase complex | 8.70E-05 |
| 1 | 11:496158 | syn | SAP190 | phosphatase complex | 8.70E-05 |
| 2 | 12:1020142 | mis | SIR3 | gene silencing | 8.70E-05 |
| 2 | 12:1020245 | mis | SIR3 | gene silencing | 8.70E-05 |
| 2 | 04:1327584 | mis | CYM1 | metalloproteaase | 8.90E-05 |
| 2 | 06:244932 | mis | BNA6 | NAD biosynthesis | 8.90E-05 |
| 1 | 05:459953 | mis | UBP5* | ubiquitin-dep. protease | 9.20E-05 |
| 3 | 13:599732 | mis | ALD3* | aldose reductase | 9.40E-05 |
| 3 | 15:840149 | syn | RIM20 | transcription reglugator | 9.60E-05 |
| 2 | 05:24902 | int | HXT13 / | hexose transporter | 9.60E-05 |
| 2 | 05:24902 | int | YEL068C | uncharacterized | 9.60E-05 |
| 4 | 15:983117 | syn | REV1* | DNA damage repair | 9.90E-05 |

**Fig 3.**
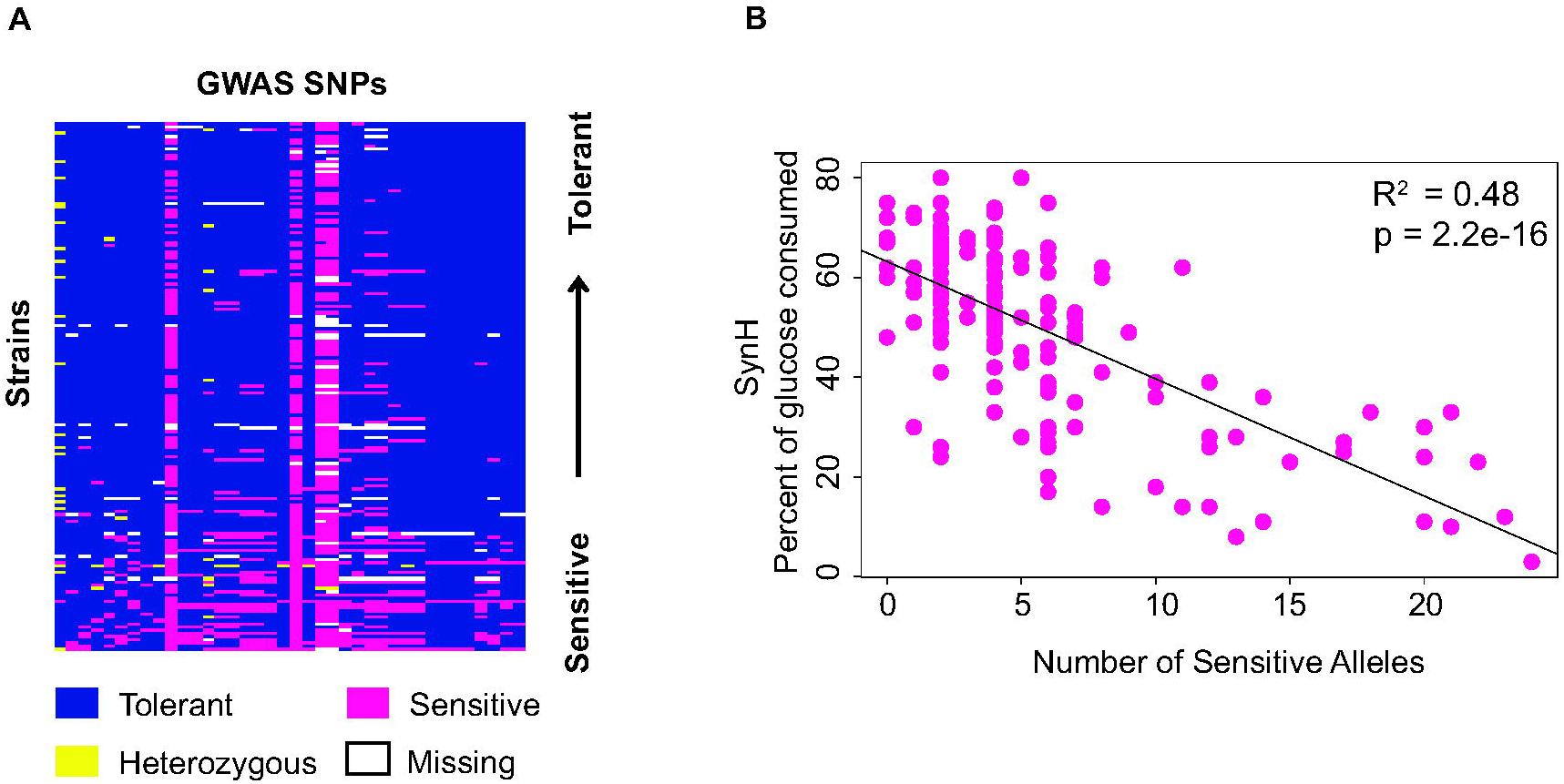
Distribution of SNP alleles. (A) A heat map of the 38 SNPs found in the GWA analysis (columns) in each strain (rows), where the alleles associated with the sensitive or resistance phenotypes are color-coded according to the key. Strains were organized from tolerant (top) to sensitive (bottom). (B) Percent glucose consumed in SynH + HTs was plotted against the number of sensitive alleles identified in each strain. Correlation of the two is indicated by the R^2^ and linear fit line.

Interestingly, the genes associated with the 38 implicated SNPs capture functionally related processes, suggesting mechanistic underpinnings of hydrolysate tolerance. Lignocellulosic hydrolysate contains a large number of toxins that affect multiple cellular functions and can target energy stores, membrane fluidity, protein and DNA integrity, and other processes [10, 65]. Our analysis implicated several genes involved in redox reactions *(ADH4, ALD3)*, protein folding or modification *(CYM1, UBP5, UFD2, AOS1)*, ergosterol or fatty acid synthesis *(ERG12, HMG2, NSG2)*, DNA metabolism and repair *(REV1, DAT1, MCM5, SHE1)*, mRNA transcription and export *(LEU3, SIR3, ELF1, RIM20, MEX67)*, mitochondrial function *(MNE1, MAS1)*, and flocculation *(FLO1, FLO10)*. Several of these processes were already known to be associated with hydrolysate stress, including flocculation [71], ubiquitin-dependent processes that may be linked to protein folding challenges [13, 72, 73], and sterol biosynthesis which affects tolerance to multiple stresses present in this media [32, 74, 75]. Nearly a third of these genes were identified as differentially expressed in our previous study comparing strain responses to SynH and rich medium [32], although this was not enriched above what is expected by chance. Thus, although gene expression differences can be informative in suggesting affected cellular processes, many of the genes implicated by GWA cannot be predicted by expression differences, especially SNPs that affect function without altering gene expression. Additional genes identified here belong to functional groups previously identified in our differential expression analysis, such as amino-acid and NAD biosynthesis.

### Gene knockouts confirm functional requirement for some implicated genes

We sought to confirm the importance of the GWA-implicated genes in SynH tolerance, first through gene-knockout analysis and then with allelic replacement in two different strains backgrounds. We began by knocking out 19 of the implicated genes in the tolerant North American strain, YPS128. Of these, 37% (7/19) of the knockout mutants had a significant phenotype when grown on SynH: four displayed decreased SynH tolerance, while 3 showed increased performance (Fig 4A). We note that 4 of the 7 knockouts had a mild phenotypic effect in standard growth medium (that was generally exacerbated in SynH), while 3 of these had a phenotype only in response to SynH (S4 Fig). The most significant knockouts decreasing tolerance in the YPS128 strain included the transcription factor *LEU3*, ribosomal protein *RPL21B*, protein phosphatase subunit *SAP190*, and to a milder extend the mitotic spindle protein *SHE1*. None of these genes has been directly implicated in tolerance to hydrolysate in previous studies.

**Fig 4.**
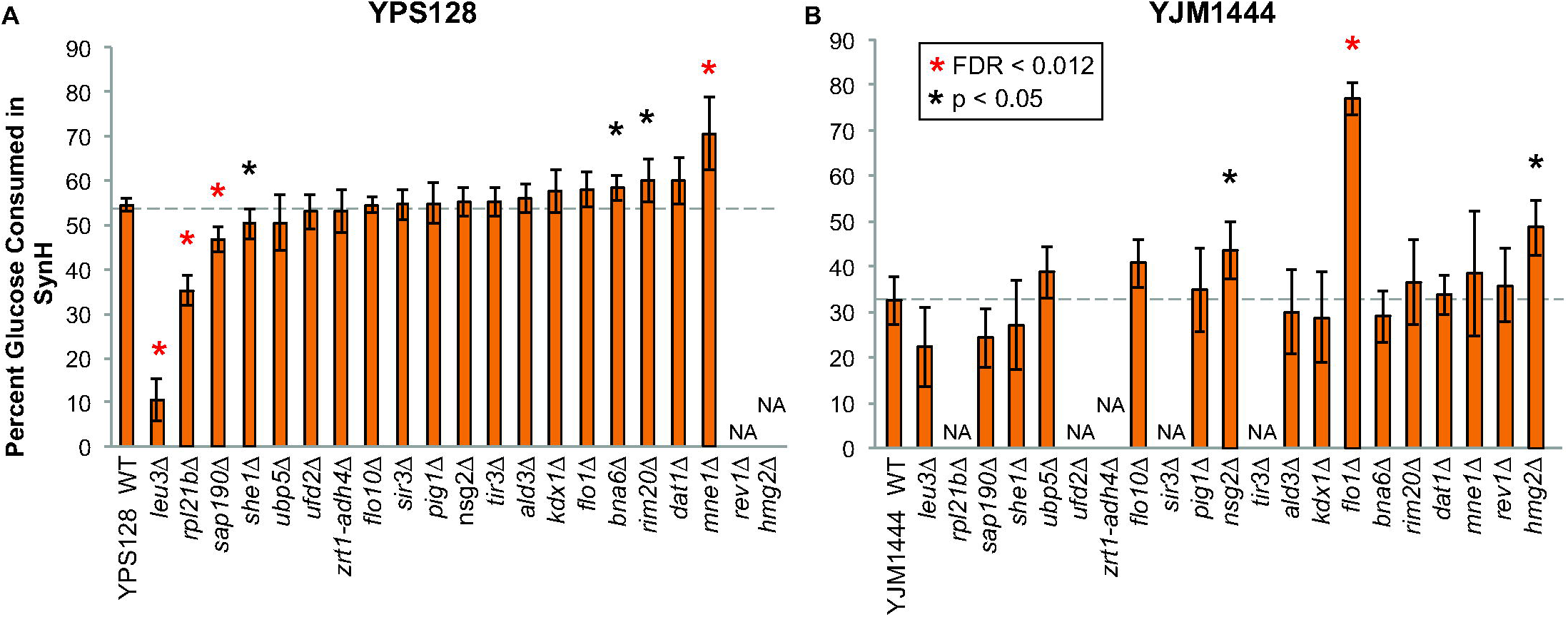
Knockout effects of genes containing SNPs found in GWA. Genes linked to SNPs implicated by GWA were deleted in one or two genetic backgrounds, tolerant strain YPS128 (A) and sensitive strain YJM1444 (B). Significance was determined by paired T-test (where experiments were paired by replicate date, see Methods) with FDR correction compared to respective wild type strain. Asterisks indicate FDR < 0.05 or p< 0.05 (which corresponds to FDR of ~13%), according to the key. Deletion strains in (B) are ordered as in (A); NA indicates missing data due our inability to make the gene deletion in that background. *zrt1-adh4Δ* indicates the deletion of an intergenic sequence between these genes.

The effect of deleting *LEU3*, which encodes the leucine-responsive transcription factor, was intriguing, since our prior work reported that amino-acid biosynthesis genes are induced specifically in response to HTs [32]. To confirm that this response was due to the toxicity found in the media and not due to amino acid shortage in SynH, we compared growth in synthetic complete (SC) medium, which has similar levels of branched-chain amino acids compared to SynH. The *LEU3* knockout strain grew as well as the wild type in SC, but it grew to 54% lower final density in SynH–HT medium and 79% lower density in SynH medium with the toxins added (Fig 5A). The defect was not fully complemented by supplementing synthetic hydrolysate with 10X the normal amino acid mix (Fig 5B), indicating that amino acid shortage in the medium is unlikely to fully explain the growth defect.

**Fig 5.**
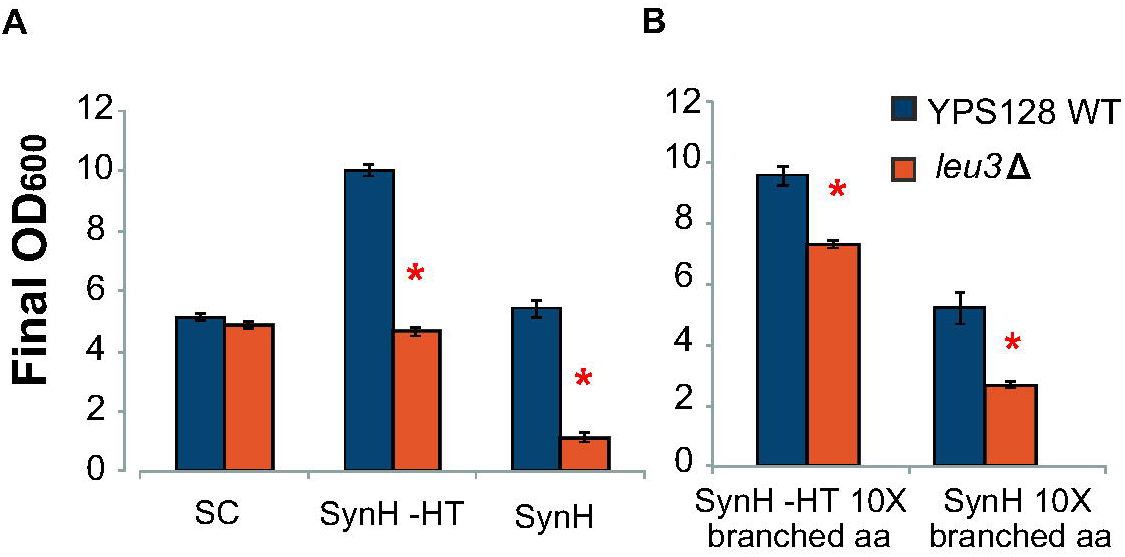
*LEU3* is important for SynH tolerance. (A) Wild-type YPS128 and a YPS128 *leu3Δ* mutant were grown in Synthetic Complete medium (SC), SynH without toxins (SynH–HT), or SynH with toxins, and final OD_600_ was measured after 24 hours. (B) Final OD_600_ was also measured in strains grown in media with 10X SC concentration of branched amino acids (leucine, isoleucine, and valine) in SynH–HT and SynH. Data represent average of 3 replicates with standard deviation. Significance was determined by paired t-test.

The most striking phenotypic improvement was caused by deletion *MNE1*, encoding a splicing factor for the cytochrome c oxidase-encoding *COX1* mRNA [76]. Aerobically, the mutant grew to roughly similar cell densities but consumed 44.7% more glucose and generated 64% more ethanol than the wild type, generating significantly more ethanol per cell (S5 Fig). A logical hypothesis is that this mutant has a defect in respiration and thus relies more on glycolysis to generate ATP and ethanol than wild-type cells [76]. Under this hypothesis, the effect of the mutation should be normalized when cells are grown anaerobically because both the mutant and wild type must rely on fermentation. However, under anaerobic conditions the mutant grew significantly better than the wild type (Fig 6A), consumed 70% more glucose (Fig 6B), and produced 63% more ethanol after 24-hour growth (Fig 6C). Thus, a simple defect in respiration is unlikely to explain the result, suggesting that Mne 1 may have a separable role relating anaerobic toxin tolerance and/or metabolism.

**Fig 6.**
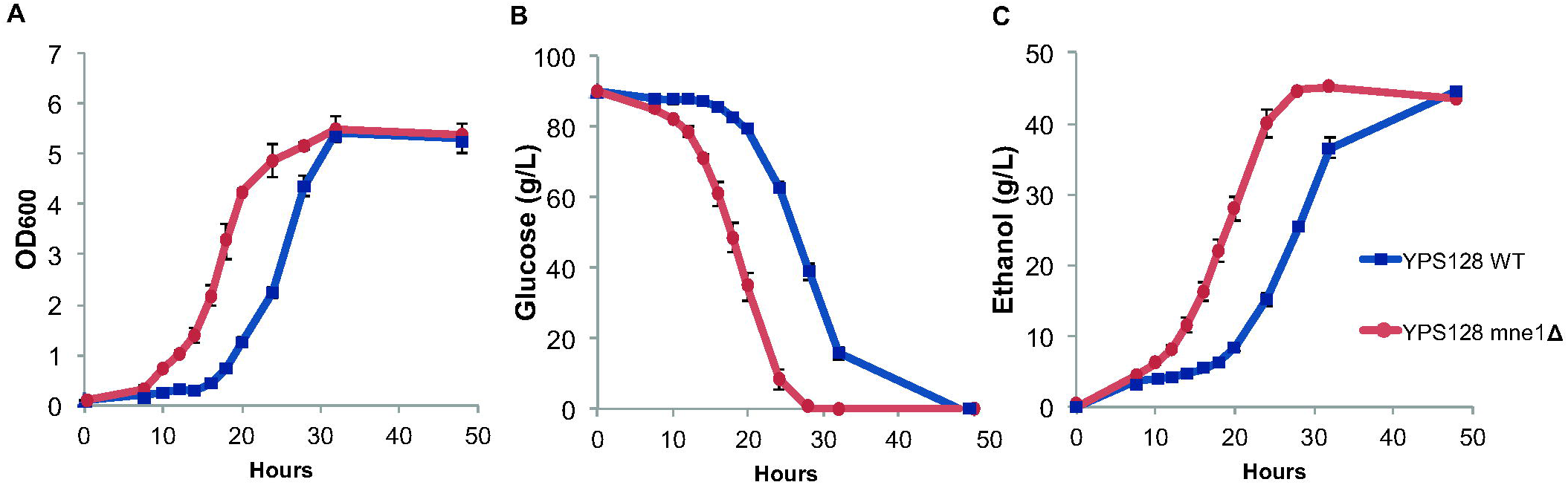
Increased SynH performance in the *mne1△* mutant is independent of oxygen availability. Wild type YPS128 and the *mne1△* mutant were grown anaerobically as described in Methods, and media was sampled over time to determine (A) cell density, (B) glucose consumption, and (C) ethanol production over time. Plots represent the average and standard deviation of 3 replicates.

### Extensive background effects influence gene involvement in SynH tolerance

We next knocked out 16 genes in the sensitive strain YJM1444, with the intention of allelic exchange (Fig 4B). We were unable to recover knockouts for some of the genes tested in YPS128, but of those we acquired 14 overlapped the YPS128 knockouts, and two *(REV2* and *HMG2)* that we were unable to knock out in the tolerant strain were added. Remarkably, knockouts had strikingly different effects between the two genetic backgrounds - while three of the gene deletions affected hydrolysate tolerance in YJM1444, there was no overlap with the gene deletions causing a statistically significant effect in YPS128 (although some mild effects may be below our statistical power to detect). The three knockouts specific to YJM1444 improved SynH tolerance and included two genes involved in sterol biosynthesis *(NSG2* and *HMG2)* and one involved in flocculation (Fig 4B). In fact, deletion of *FLO1* dramatically reduced the flocculation phenotype of YJM1444 and resulted in >236% increased glucose consumption in SynH. This single mutation converted YJM1444 tolerance to the level of SynH tolerance seen in YPS128 (S6 Fig). To test that this phenotypic effect was directly caused by the *FLO1* allele, we deleted its paralog *FLO5*, which caused neither a change in flocculation nor increased glucose consumption of the culture (S7 Fig).

There appeared to be subtle, but not significant, effects of the *MNE1* deletion in YJM1444 and we wondered if the was obscured by flocculation. Therefore, we measured glucose consumption in high-rpm shake flasks that disrupt flocculation. Indeed, *MNE1* deletion had a significant benefit under these conditions; however, the magnitude of the effect was more subtle than *MNE1* deletion in YPS128 (S8A Fig). We also tested this deletion in an industrial strain, Ethanol Red (E. Red). Deletion of *MNE1* in a haploid spore derived from E. Red produced a minor, reproducible benefit although it was not statistically significant (S8A Fig). Nonetheless, these results show that *MNE1* plays a role in SynH tolerant, albeit to different levels, in three different strain backgrounds.

### Genetic background effects dominate the effect of allelic variation in HT tolerance

We tested allelic effects in two ways. First, we introduced a plasmid-borne copy of the tolerant allele or sensitive allele (S5 Table) into YPS128 lacking the native gene, and measured percent final glucose consumption in SynH (S9 Fig) in synthetic complete medium (required to allow drug-based plasmid selection) with HTs (Fig 7A). The assay was fairly noisy, in part because wild-strains are somewhat intolerant to the plasmid drug marker (unpublished). Nonetheless, there was a clear effect for the *FLO1* allele, which caused YPS128 to become flocculant and dramatically decreased growth in the SC with HTs. We did not observe other allele-dependent effects that overcame the variability of the assay, including for the genes whose knockout produced a defect in YPS128. Second, we performed reciprocal hemizygosity analysis for six genes, including three genes that whose deletion produced differential effects in YPS128 and YJM144. We crossed the YPS128 and YJM1444 backgrounds such that the resulting diploid was hemizigous for either the tolerant or sensitive allele (Fig 7B). In this case, none of the six genes had an allele-specific effect - surprisingly, this included *FLO1* for which there was clear allelic impact in the haploid backgrounds. We realized a unique phenotype in the YPS128-YJM1444 hybrid: whereas the strain is heterozygous for the functional *FLO1* allele, the hybrid lost much of the flocculence of the YJM1444 strain (S8B Fig). *FLO1* expression is known to be repressed in some diploid strains [77]. Thus, simply mating the strains in effect created a new genetic background that changed the allelic impact of the gene.

**Fig 7.**
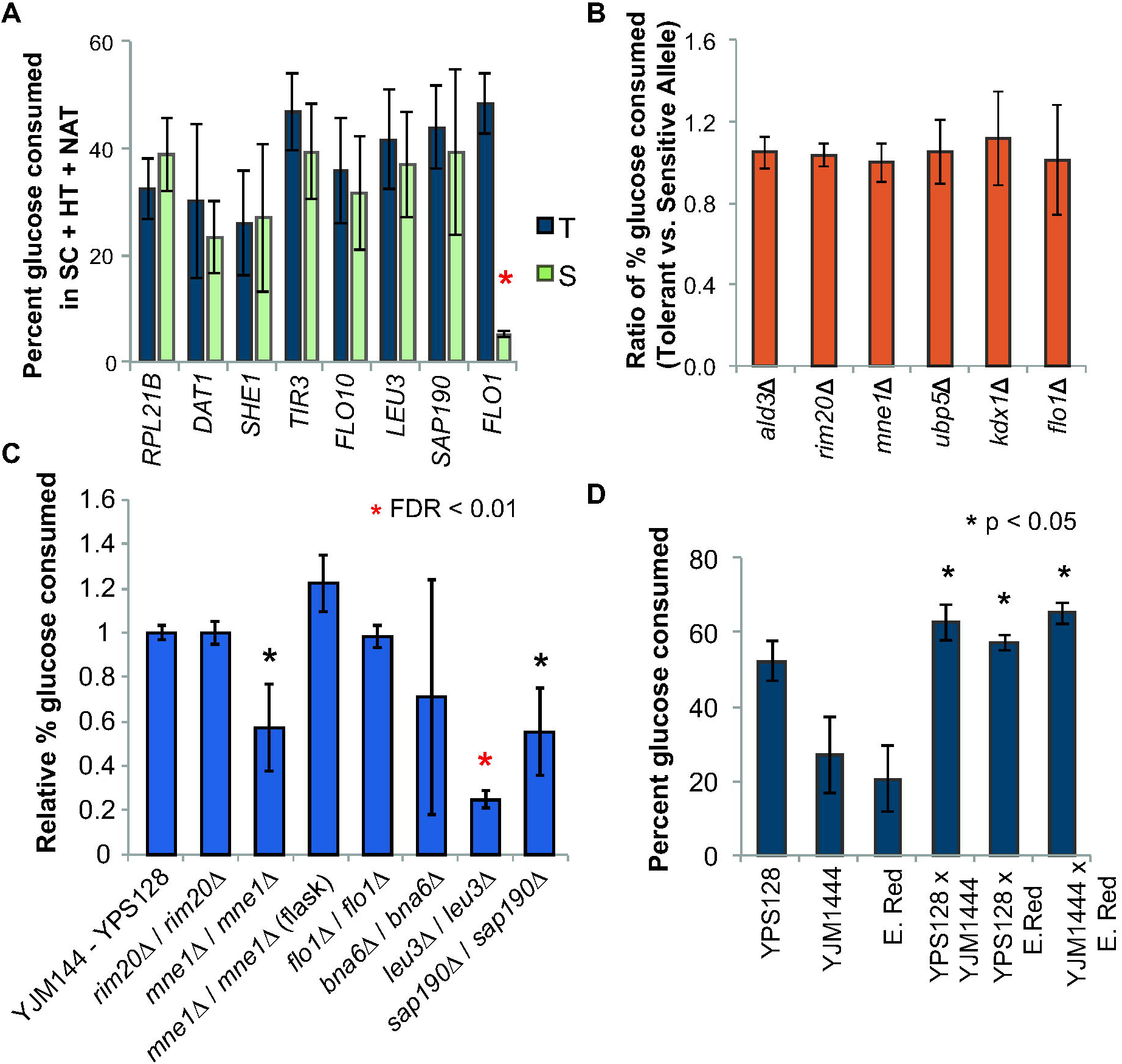
Little allele-specific contribution to SynH tolerance. (A) YPS128 strains lacking individual genes were complemented with a plasmid carrying the tolerant allele (T) or the sensitive allele (S) of each gene. Cells were grown in synthetic complete medium (SC) with HTs and nourseothricin (NAT) to allow for drug-based plasmid selection. Significance was determined by paired t-test comparing strains carrying the tolerant versus sensitive allele. Data represent the average and standard deviation of three replicates. (B) Relative phenotype based on reciprocal hemizygosity analysis (RHA) where the ratio of glucose consumption of strains carrying the tolerant versus sensitive allele was calculated across 7 biological replicates. (C) Relative percent glucose consumed in wild-type YJM144 x YPS128 hybrid and homozygous deletion strains compared to the average of the wild-type YJM144 x YPS128 hybrids, in SynH as described in Methods. Data represent the average and standard deviation of three replicates. One of the *bna6A* cultures did not grow; the other two replicates looked indistinguishable from the wild-type culture. (D) Phenotypic improvement achieved by crossing strains with diverse phenotypic and genetic characteristics was investigated by measuring percent of glucose consumption in SynH. Significance was determined by paired t-test comparing each hybrid and the most tolerant strain, YPS128.

We wondered if this effect explained the lack of allele-specific phenotypes for other implicated genes. We therefore created homozygous deletions in the diploid hybrid for six genes whose deletion had strain-specific impacts in the haploids (Fig 7C). Two of the knockouts *(leu3Δ* and *sap190Δ*) produced a defect in the hybrid, similar to the effect seen in YPS128. Homozygous deletion of *MNE1* produced a unique growth defect in 24-well plates that was not seen in the haploids or the hemizigous diploids. This appeared to be due to increased flocculation in the hybrid diploid; growth in shake flasks to disrupt flocculation resulted in a mild but statistically insignificant benefit to the hybrid when grown in flasks, similar to that seen for YJM1444. In contrast, deletion of *RIM20* or *FLO1* had no effect under these growth conditions - this explains the lack of allele specific effect, because the genes are no longer important in this background and under these growth conditions.

Mating YJM128 and YJM144 created a new background that surpassed performance of YPS128 (Fig 7D). We wondered if hybridization could benefit other strains as well. We mated industrial strain E. Red crossed to YJM1444 and YPS128. E. Red and YJM1444 were both scored as sensitive and perform similarly in SynH (Fig 7D). However, the hybrid had a striking jump in SynH tolerance, exceeding the tolerance of YPS128. This benefit may be in part because the new diploid background changes the flocculation phenotype. On the other hand, YJM144 and E. Red harbor alternate alleles at 71% of the SNPs implicated by GWA, raising the possibility that complementation of recessive alleles could also contribute to the strain improvement (see Discussion).

## Discussion

Engineering strains for tolerance to lignocellulosic hydrolysate has proven difficult due to the complex stress responses required to deal with the combinatory effects of toxins, high osmolarity, and end products such as alcohols and other chemicals. Even when the cellular targets of stressors are known, the mechanisms for increasing tolerance are not always clear. We leveraged phenotypic and genetic variation to implicate new mechanisms of hydrolysate tolerance, by finding correlations between phenotypic and genetic differences among a collection of *Saccharomyces cerevisiae* strains, which allowed us to implicate specific genes and alleles involved in hydrolysate tolerance. The results indicate several important points relevant to engineering improved hydrolysate tolerance and genetic architecture of tolerance more broadly.

Perhaps the most striking result is the level to which gene involvement varies across the strains in our study. We expected that swapping alleles of implicated SNPs should contribute to variation in the phenotype. Most alleles did not detectably affect tolerance, although it is likely that they may have a minor contribution below our limit of detection. Indeed, strains that harbor more deleterious alleles are significantly more sensitive to SynH (Fig 3B), as expected for an additive trait. But at the same time, we uncovered significant variation in whether the underlying gene was involved in the phenotype. Among the genes that we were able to knockout in both strains (14 genes), 57% produced a phenotype (to varying levels and significance) in one of the two strains we tested. This indicates substantial epistatic interactions with the genetic background, such that the gene is important in one strain and but dispensable in another. Even more striking is the case of *FLO1:* knocking out the functional gene in YJM1444 produced a major benefit to that strain, whereas introducing the functional allele to YPS128 was very detrimental to SynH tolerance. Yet neither the allele nor the gene itself influenced SynH tolerance in the hybrid, because the hybrid is much less flocculant under these conditions (despite carrying functional YJM1444 *FLO1* gene).

While it may not be surprising that gene knockouts result in quantitatively different phenotypes, we did not expect that most knockouts would have no detectible effect in specific backgrounds. It will be important to investigate the extent to which this effect is true in other organisms and for other phenotypes. However, evidence in the literature hints at the breadth of this result: several genes are required for viability in one yeast strain but not another [63, 78], while overexpression of other genes produces a phenotype in one background but not others [32]. Genetic background effects on gene contributions have been reported before, in yeast and other organisms [35, 79–84]; however, the extent to which different genes appear to be involved in toxin tolerance in the different strains studied here suggests an important consideration that is underappreciated in GWA analysis: that the network of genes contributing to the phenotype could be largely different depending on genomic context. Dissecting these epistatic interactions is likely to be daunting, since a major challenge in most GWA studies remains identifying the epistatic interactions due to the high statistical hurdle [34, 85, 86]. We propose that emerging network-based approaches to augment linear contributions will be an important area in identifying genetic contributions in the context of background-specific effects.

QTL mapping has allowed the characterization of the genetic architecture of industrially relevant stresses, including tolerance to ethanol [22, 87], acetic acid [23, 56], and plant hydrolysate [25] among many others [24, 88–90]. But while this method exploits the genetic diversity between two strains, with GWA we were able to study a much larger collection of genetic diversity, providing unique insights. SynH tolerance is clearly a complex trait, with many genes likely contributing. Previous studies have shown that part of the growth inhibition can be explained by a re-routing of resources to convert toxins into less inhibitory compounds [18, 19, 91–94] and to repair damage generated by reactive oxygen species in membranes and proteins [13, 14, 95]. One of the most significant effects was caused by deletion of *LEU3*, the transcription factor regulating genes involved in branched amino acid biosynthesis. Interestingly, weak acids have been shown to inhibit uptake of aromatic amino acids causing growth arrest [96], and it is possible that Leu3 is required to combat this effect. Chemical genomic experiments suggest an additional role for Leu3 in managing oxidative stress in the cell [97], which could relate to oxidative stressors in hydrolysate [13, 14, 32]. We also uncovered a gene, Mne1, that when deleted significantly increases ethanol production in SynH. Mne1 aids the splicing of *COX1* mRNA [76] and has not been previously linked to stress tolerance. Interestingly, *MNE1* mutants produced more ethanol per cell aerobically, but also grew substantially better in SynH anaerobically, raising the possibility that Mne1 plays an additional, unknown role in cellular physiology that can be utilized to increase fermentation yields. Finally, although flocculation has been previously shown to increase cell survival in hydrolysate [71], our study showed that flocculation reduced the rate of sugar consumption in the culture, likely because cells in the middle of the clump are nutrient restricted. Together, these results shed new light on SynH tolerance and mechanisms for future engineering.

Our results raise broader implications for strain engineering, based on the genetic architectures uncovered here. Given the implication of gene-by-background interactions, the best route for improving strain performance may be crossing strains for hybrid vigor [98–100]. Indeed, we unexpectedly generated a strain that outperformed the tolerant YPS128, by crossing two poor performers in SynH. This improved vigor could emerge if the hybrid complements recessive deleterious alleles in each strain, or if mating creates a new genetic background that changes the requirements (and fitness) of the strain. We believe that both models - weak but additive allelic contributions in the context of epistatic background effects - are at work in our study. For additive traits, GWA and genomic studies can have significant practical power, by predicting where individual strains fall on the genotype-phenotype spectrum and by suggesting which strains should be crossed for maximal phenotypic effect.

## Methods

### Strains

Strains used in the GWA are listed in S1 Table Gene knockouts were performed in strains derived from North American strain YPS128 and mosaic strain YJM1444. The homozygous diploid parental strains were first engineered into stable haploids by knocking out the homothallic switching endonuclease (HO) locus with the *KAN-MX* antibiotic marker [101], followed by sporulation in 1% potassium acetate plates and dissection of tetrads to attain heterothallic MATa and MATα derivatives. Gene knockouts were generated through homologous recombination with the HERP1.1 drug resistance cassette [102] and verified by 3 or 4 diagnostic PCRs (validating that the cassette was integrated into the correct locus and that no PCR product was generated from within the gene that was deleted). Most knockouts removed the gene from ATG to stop codon, but in some cases (e.g. *kdx1)* additional flanking sequence was removed, without removing neighboring genes. Genes from YPS128 or strains carrying the sensitive allele (S5 Table) were cloned by homologous recombination onto a CEN plasmid, taking approximately 1,000 bps upstream and 600 bps downstream from each genome, and verified by diagnostic PCR. Phenotyping of strains harboring alternate alleles on plasmids was performed in as previously described, except that the pre-culture was grown in YPD with 100 mg/L nourseothricin (Werner BioAgents, Germany) to maintain the plasmid expressing each allele. We note that plasmid-bourn expression of the gene complemented the gene-deletion phenotype, where applicable, in all cases tested (not shown). Allele specific effects were additionally tested by reciprocal hemizygosity analysis (RHA) [103]. The HO locus was replaced with the nourseothricin resistance cassette *(NAT-MX)* for each mating type of YPS128 and YJM1444. These were then crossed with the appropriate deletion strain of opposite mating type and harboring the KanMX cassette, selecting for mated cells resistant to both drugs, to generate heterozygous strains that were hemizigous for the gene in question (crosses shown in S6 Table).

### Media, growth, and phenotyping conditions

Synthetic Hydrolysate (SynH) medium mimics the lignocellulosic hydrolysate generated from AFEX ammonium treated corn stover with 90 g glucan/L loading and was prepared as in Sardi *et al*. (2016). Two versions were prepared to represent the complete hydrolysate (SynH) and the hydrolysate without the hydrolysate toxin cocktail (HT) (SynH - HT), as previously published [32].

Phenotyping for GWA, gene deletion assessment, and RHA, was performed using high throughput growth assays in 24 well plates (TPP^®^ tissue culture plates, Sigma-Aldrich, St. Louis, MO). To prepare the cultures, 10 μl of thawed frozen cell stock were pinned onto YPD agar plates (1% yeast extract, 2% peptone, 2% dextrose, 2% agar) and grown for 3 days at 30°C. Cells were then pre-cultured in 24 well plates containing 1.5 ml of YPD liquid, sealed with breathable tape (AeraSeal, Sigma-Aldrich, St. Louis, MO), covered with a lid and incubated at 30°C while shaking for 24 h. Next, 10 μl of saturated culture was transferred to a 24 well plate containing 1.5 ml of SynH or SynH-HT where indicated, and grown as the preculture for 24 h. Cell density was measured by optical density at 600 nm (OD_600_) as ‘final OD’. Culture medium collected after cells were removed by centrifugation was used to determine glucose concentrations by YSI 2700 Select high performance liquid chromatography (HPLC) and refractive index detection (RID) (YSI Incorporated, Yellow Springs, OH). Biological replicates were performed on different days.

For GWA, we used four different but related phenotype measures of cells growing in SynH or SynH - HTs: 1) final OD_600_ as a measure of growth, 2) percent of starting glucose consumed after 24 hours in SynH, 3) HT tolerance based on OD_600_ (calculated as the ratio of final OD_600_ in SynH versus final OD_600_ in SynH–HTs), and 4) HT tolerance based on glucose consumption (calculated as the ratio of glucose consumed in SynH versus in SynH–HTs). Strains and phenotype scores are listed in S1 Table. Initial phenotyping for GWA was performed in biological duplicates; knockout strains and hemizigous strains were phenotyped in five biological replicates to increase statistical power, whereas homozygous deletion strains were phenotyped in triplicate. Replicates for each batch of strains shown in each figure were performed on separate days, for paired statistical analysis.

Experiments done for allele replacements expressed on plasmids were performed in glass tubes using modified synthetic complete medium (SC) with high sugar concentrations and the toxin cocktail where indicated (Sardi *et al* 2016) to mimic SynH but with no ammonium to support nourseothricin selection [104] (1.7 g/L YNB w/o ammonia sulfate and amino acids, 1 g/L monosodium glutamic acid, 2 g/L amino acid drop-out lacking leucine, 48 μg/L leucine, 90 g/L dextrose, 45 g/L xylose). This was required since nourseothricin selection does not work in high-ammonium containing SynH. First, we precultured strains carrying plasmids in SC medium with nourseothricin (200 ug/ml) for 24 h. Next, we inoculated a fresh culture at a starting OD_600_ of 0.1 in 7 ml of the modified synthetic complete medium with nourseothricin (200 ug/ml) and HTs. Cultures were grown for 24 h and phenotyped as described above. Replicates were performed on different days, and thus samples were paired by replicate date for t-test analysis.

Anaerobic phenotyping was performed in the anaerobic chamber, where cells were grown in flasks containing 25 ml SynH or SynH-HT and maintained in suspension using a magnetic stir bar. Ethanol production was measured over time by HPLC RID analysis. Paired t-test analysis was performed to determine significance, pairing samples by replicate date.

### Genomic sequencing and Analysis

We obtained publicly available whole genome sequencing reads from *Saccharomyces cerevisiae* sequencing projects [39, 42, 52, 60]. Sequencing reads were mapped to reference genome S288C (NC_001133, version 64 [105]) using bwa-mem [106] with default settings. Single nucleotide polymorphisms (SNPs) were identified using GATK [107] Unified Genotyper, analyzing all the strains together to increase detection power. GATK pipeline included base quality score calibration, indel realignment, duplicate removal, and depth coverage analysis. Default parameters were used except for -mbq 25 to reduce false positives. Variants were filtered using GATK suggested criteria: QD < 2, FS > 60, MQ < 40. A dataset with high quality SNPs was generated using VCFtools [108] by applying additional filters of a quality value above 2000 and excluding sites with more that 80% missing data. Genetic variant annotation was performed using SNPEff [109]. Principal component analysis and the neighbor-joining tree were performed with the R package Adegenet 1.3–1 [110] using the entire collection of high quality SNPs (486,302 SNPs).

### Genome-wide association analysis

Correlations between genotype and phenotype were performed using a mixed linear model implemented in the software GAPIT [68]. Only SNPs with a minor allele frequency (MAF) of at least 2% were used for this analysis (282,150 SNPs). Multiple models, each incorporating a different number of principal components to capture population structure (from 0 - 3), were analyzed. The final model was manually chosen as the one with the greatest overall agreement between the distribution of expected and the observed p-values, *i.e*. based on QQ plots with the least skew across the majority of SNPs. We performed four analyses, one for each for the four related phenotypes measured. The model used to map SynH final OD_600_ and SynH percent glucose consumed used 0 principal components, with population structure corrected using only the kinship generated by GAPIT. The model used to map HT tolerance based on relative final OD_600_ used 2 principal components, and the model to map HT tolerance based on glucose consumed incorporated 1 principal component. The threshold for significance accounting for multiple-test correction was identified by dividing the critical p-value cutoff of 0.05 by the number of independent tests estimated by the SimpleM method [111], which decreased the number of tests from 282,150 to 137,398 to produce a p-value threshold of 3.6e-7 [112]. However, none of our tests passed this threshold, which is likely overly conservative. We therefore used a p-value threshold of 1e-04 to identify genes for detailed follow-up analysis. We realized that the extreme phenotypes of Asian/sake strains coupled with their strong population structure might be confounding the analysis (unpublished). Therefore, to further reduce the chance of false positives due to residual population influences, we reran the analyses without the 11 sake strains and removed from the original list of significant SNPs those with p>5e-3. For each locus carrying a significant SNP, we plotted phenotypic distributions for each possible genotype. We focused subsequent downstream analysis on individual SNPs whose effects were additive across strains that were heterozygous and homozygous at that site, assessed visually. Genes affected by each SNP were determined by the SNPEff annotation, which predicted the effect of variants on genes.

## Acknowledgments

We thank Y. Zhang for generating SynH medium and T. Sato for helpful comments on this work. This work was supported by the Department of Energy Great Lakes Bioenergy Research Center (DOE Office of Science BER DE-FC02-07ER64494). MS was supported by a National Science Foundation Graduate Research Fellowship (DGE-1256259).

## Supporting information

**S1 Figure. Strain-specific differences in SynH and HT tolerance.** Tolerance to lignocellulosic hydrolysate across strains (left) and summarized within each population (right) measured as final OD_600_ in SynH (A) and HT tolerance (B) (based the ratio of final OD_600_ in cultures grown with åversus without toxins). Strains were ordered as in (A). Population names for boxplots are listed in Fig 1.

**S2 Figure. Allele frequency of significant SNPs found through GWA.** Distribution of the minor allele frequency of 76 SNPs is shown.

**S3 Figure. Distribution of tolerant and sensitive allele based on population.** A heat map of the 38 SNPs found in the GWA analysis (columns) in each strain (rows), where the alleles associated with the sensitive or resistance phenotypes are color-coded according to the key. Strains were grouped based on their ancestral population as indicated in the figure; Wine, Asian, NA (North American), WA (West African), and MOS (mosaic).

**S4 Figure. Knockout effects of genes containing SNPs found in GWA when cells were grown in rich medium.** The phenotypic impact of genes affected by SNPs found in GWA was tested in rich lab medium (YPD). Average and standard deviation of 5 replicates is shown. Significance was determined by paired t-test compared to wild type strain.

**S5 Figure. Deletion of *MNE1* significantly increases fermentation rates in SynH.** Effects of the *MNE1* deletion in YPS128 were measured in cells growing in flasks. We observed increased glucose consumption (A), higher production of ethanol (B), and higher production of ethanol per cell (C). Average and standard deviation of 3 replicates is shown. Significance was determined by paired t-test compared to wild type strain.

**S6 Figure. Deletion of *FLO1* significantly increases YJM1444 glucose consumption in SynH.** Effects of the *FLO1* deletion in YPS128 and YJM1444 were measured in cells growing in flasks. This single deletion increased YJM1444 glucose consumption in SynH to the level seen in YPS128. Significance was determined by paired t-test compared to wild type strain.

**S7 Figure. Increased tolerance to SynH is specific to the *FLO1* deletion.** Deletion of *FLO5*, the paralog of *FLO1*, did not decrease flocculation (not shown) or increase growth in SynH.

**S8 Figure. Deletion of *MNE1* improves glucose consumption in SynH in multiple genetic backgrounds. (A)** Effect of *MNE1* deletion on glucose consumption in SynH was measured in YJM1444 and the ethanol red strain (E. Red) by growing cells in flasks and measuring percent of glucose consumed after 24 hours. Significance was determined by paired t-test compared to wild type strain. Red asterisk symbolizes P < 0.01. **(B)** Flocculation differences in haploid strains (left), and three independently made crosses of YPS128 and YJM1444 from the designated mating types. Cultures were grown in tubes to saturation and cells allowed to sit briefly without shaking. These culture conditions exacerbate the amount of flocculation for easy visualization. The YJM1444 haploid is highlight flocculant under these conditions (visualized as clear media with cell precipitate at the bottom of the tube) whereas multiple independently made hybrids are no longer flocculant.

**S9 Figure. Plasmid complementation carrying tolerant and sensitive allele in SynH.** YPS128 deletion mutants were transformed with an empty plasmid (pKI), a plasmid carrying the tolerant allele (pT), or a plasmid carrying the sensitive allele (pS). Cells were grown in SynH, which does not allow the use antibiotics due to the presence of ammonium sulfate in the medium. Although most cells likely retain the plasmid over the duration of this experiment, we were unable to detect allele-specific effects that overcome the variation in the experiments.

**S1 Table. Strain information**

**S2 Table. Summary of SNPs and predicted impacts.** SNP classifications were performed by SnpEff as outlined in Methods. Low impact genic polymorphisms are represented by synonymous codon changes, moderate impact genic SNPs are nonsynonymous codon changes, and high impact variants include introduction of premature stop codons, altered start position, or interruptions of slicing regions.

**S3 Table. List of genes with high impact mutations.**

**S4 Table. Initial identification of SNPs correlated with SynH tolerance.** Initial set of SNPs whose p-value passed our threshold in any of the GWA are shown, ranked by significance. Phenotypes to which the SNP was associated are listed in the first column; (1) Final OD_600_ in SynH, (2) Percent of glucose consumed in SynH, (3) HT tolerance based on OD_600_, (4) HT tolerance based on glucose consumed. SNPs identified in multiple GWA, the most significant p-value is listed in the last column. SNP type was determined by SNPeff: syn, synonymous; mis, missense; int, intergenic.

**S5 Table. Plasmids with tolerant and sensitive alleles.** Strain genotype sources for cloning of tolerant and sensitive allele for plasmid complementation are shown.

**S6 Table. Strains used for RHA between YPS128 and YJM1444 to test alleles found in GWAS.** The type of allele that is tested in each hybrid is labeled as sensitive (S) and tolerant (T).

## References

1. Fairley P. Introduction: Next generation biofuels. Nature. 2011;474(7352):S2–5. doi: 10.1038/474S02a. PubMed PMID: 21697838.

2. Perez J, Munoz-Dorado J, de la Rubia T, Martinez J. Biodegradation and biological treatments of cellulose, hemicellulose and lignin: an overview. Int Microbiol. 2002;5(2):53–63. doi: 10.1007/s10123-002-0062-3. PubMed PMID: 12180781.

3. Balan V, Bals B, Chundawat SP, Marshall D, Dale BE. Lignocellulosic biomass pretreatment using AFEX. Methods Mol Biol. 2009;581. doi: 10.1007/978-1-60761-214-8_5.

4. Hendriks AT, Zeeman G. Pretreatments to enhance the digestibility of lignocellulosic biomass. Bioresour Technol. 2009;100(1):10–8. doi: 10.1016/j.biortech.2008.05.027. PubMed PMID: 18599291.

5. Chundawat SP, Beckham GT, Himmel ME, Dale BE. Deconstruction of lignocellulosic biomass to fuels and chemicals. Annu Rev Chem Biomol Eng. 2011;2:121–45. doi: 10.1146/annurev-chembioeng-061010-114205. PubMed PMID: 22432613.

6. Stephanopoulos G. Challenges in engineering microbes for biofuels production. Science. 2007;315(5813):801–4. doi: 10.1126/science.1139612. PubMed PMID: 17289987.

7. Klinke HB, Thomsen AB, Ahring BK. Inhibition of ethanol-producing yeast and bacteria by degradation products produced during pre-treatment of biomass. Appl Microbiol Biotechnol. 2004;66(1):10–26. doi: 10.1007/s00253-004-1642-2. PubMed PMID: 15300416.

8. Larsson S, Quintana-Sainz A, Reimann A, Nilvebrant NO, Jonsson LJ. Influence of lignocellulose-derived aromatic compounds on oxygen-limited growth and ethanolic fermentation by Saccharomyces cerevisiae. Appl Biochem Biotechnol. 2000;84–86:617–32. PubMed PMID: 10849822.

9. Almeida JRM, Modig T, Petersson A, Hähn-Hägerdal B, Lidén G, Gorwa-Grauslund MF. Increased tolerance and conversion of inhibitors in lignocellulosic hydrolysates by Saccharomyces cerevisiae. Journal of Chemical Technology & Biotechnology. 2007;82(4):340–9. doi: 10.1002/jctb.1676.

10. Piotrowski JS, Zhang YP, Sato T, Ong I, Keating D, Bates D. Death by a thoU S And cuts: the challenges and diverse landscape of lignocellulosic hydrolysate inhibitors. Front Microbiol. 2014;5. doi: 10.3389/fmicb.2014.00090.

11. Chundawat SP, Vismeh R, Sharma LN, Humpula JF, da Costa Sousa L, Chambliss CK. Multifaceted characterization of cell wall decomposition products formed during ammonia fiber expansion (AFEX) and dilute acid based pretreatments. Bioresour Technol. 2010;101. doi: 10.1016/j.biortech.2010.06.027.

12. Krebs HA, Wiggins D, Stubbs M, Sols A, Bedoya F. Studies on the mechanism of the antifungal action of benzoate. Biochemical Journal. 1983;214(3):657–63. PubMed PMID: PMC1152300.

13. Nguyen TT, Iwaki A, Ohya Y, Izawa S. Vanillin causes the activation of Yap 1 and mitochondrial fragmentation in Saccharomyces cerevisiae. J Biosci Bioeng. 2014;117(1):33–8. doi: 10.1016/j.jbiosc.2013.06.008. PubMed PMID: 23850265.

14. Allen SA, Clark W, McCaffery JM, Cai Z, Lanctot A, Slininger PJ, et al. Furfural induces reactive oxygen species accumulation and cellular damage in Saccharomyces cerevisiae. Biotechnol Biofuels. 2010;3:2. doi: 10.1186/1754–6834-3-2. PubMed PMID: 20150993; PubMed Central PMCID: PMCPMC2820483.

15. Modig T, Liden G, Taherzadeh MJ. Inhibition effects of furfural on alcohol dehydrogenase, aldehyde dehydrogenase and pyruvate dehydrogenase. Biochem J. 2002;363(Pt 3):769–76. PubMed PMID: 11964178; PubMed Central PMCID: PMCPMC1222530.

16. Pisithkul T, Jacobson TB, O'Brien TJ, Stevenson DM, Amador-Noguez D. Phenolic Amides Are Potent Inhibitors of De Novo Nucleotide Biosynthesis. Appl Environ Microbiol. 2015;81(17):5761–72. doi: 10.1128/AEM.01324-15. PubMed PMID: 26070680; PubMed Central PMCID: PMCPMC4551265.

17. Iwaki A, Ohnuki S, Suga Y, Izawa S, Ohya Y. Vanillin inhibits translation and induces messenger ribonucleoprotein (mRNP) granule formation in saccharomyces cerevisiae: application and validation of high-content, image-based profiling. PLoS One. 2013;8(4):e61748. doi: 10.1371/journal.pone.0061748. PubMed PMID: 23637899; PubMed Central PMCID: PMCPMC3634847.

18. Petersson A, Almeida JR, Modig T, Karhumaa K, Hahn-Hagerdal B, Gorwa-Grauslund MF, et al. A 5-hydroxymethyl furfural reducing enzyme encoded by the Saccharomyces cerevisiae ADH6 gene conveys HMF tolerance. Yeast (Chichester, England). 2006;23(6):455–64. doi: 10.1002/yea.1370. PubMed PMID: 16652391.

19. Liu ZL, Moon J, Andersh BJ, Slininger PJ, Weber S. Multiple gene-mediated NAD(P)H-dependent aldehyde reduction is a mechanism of in situ detoxification of furfural and 5-hydroxymethylfurfural by Saccharomyces cerevisiae. Appl Microbiol Biotechnol. 2008;81(4):743–53. doi: 10.1007/s00253-008-1702-0. PubMed PMID: 18810428.

20. Teixeira MC, Raposo LR, Mira NP, Lourenco AB, Sa-Correia I. Genome-wide identification of Saccharomyces cerevisiae genes required for maximal tolerance to ethanol. Appl Environ Microbiol. 2009;75. doi: 10.1128/aem.00845-09.

21. Mira NP, Palma M, Guerreiro JF, Sa-Correia I. Genome-wide identification of Saccharomyces cerevisiae genes required for tolerance to acetic acid. Microb Cell Fact. 2010;9:79. doi: 10.1186/1475-2859-9-79. PubMed PMID: 20973990; PubMed Central PMCID: PMCPMC2972246.

22. Swinnen S, Schaerlaekens K, Pais T, Claesen J, Hubmann G, Yang Y, et al. Identification of novel causative genes determining the complex trait of high ethanol tolerance in yeast using pooled-segregant whole-genome sequence analysis. Genome Res. 2012;22(5):975–84. doi: 10.1101/gr.131698.111. PubMed PMID: 22399573; PubMed Central PMCID: PMCPMC3337442.

23. Geng P, Xiao Y, Hu Y, Sun H, Xue W, Zhang L, et al. Genetic dissection of acetic acid tolerance in Saccharomyces cerevisiae. World J Microbiol Biotechnol. 2016;32(9):145. doi: 10.1007/s11274-016-2101-9. PubMed PMID: 27430512.

24. Hubmann G, Mathe L, Foulquie-Moreno MR, Duitama J, Nevoigt E, Thevelein JM. Identification of multiple interacting alleles conferring low glycerol and high ethanol yield in Saccharomyces cerevisiae ethanolic fermentation. Biotechnol Biofuels. 2013;6(1):87. doi: 10.1186/1754-6834-6-87. PubMed PMID: 23759206; PubMed Central PMCID: PMCPMC3687583.

25. Maurer MJ, Sutardja L, Pinel D, Bauer S, Muehlbauer AL, Ames TD, et al. Quantitative Trait Loci (QTL)-Guided Metabolic Engineering of a Complex Trait. ACS Synth Biol. 2016. doi: 10.1021/acssynbio.6b00264. PubMed PMID: 27936603.

26. Chen Y, Sheng J, Jiang T, Stevens J, Feng X, Wei N. Transcriptional profiling reveals molecular basis and novel genetic targets for improved resistance to multiple fermentation inhibitors in Saccharomyces cerevisiae. Biotechnol Biofuels. 2016;9:9. doi: 10.1186/s13068-015-0418-5. PubMed PMID: 26766964; PubMed Central PMCID: PMCPMC4710983.

27. Heer D, Sauer U. Identification of furfural as a key toxin in lignocellulosic hydrolysates and evolution of a tolerant yeast strain. Microbial biotechnology. 2008; 1(6):497–506. doi: 10.1111/j.1751-7915.2008.00050.x. PubMed PMID: PMC3815291.

28. Hawkins GM, Doran-Peterson J. A strain of Saccharomyces cerevisiae evolved for fermentation of lignocellulosic biomass displays improved growth and fermentative ability in high solids concentrations and in the presence of inhibitory compounds. Biotechnol Biofuels. 2011;4(1):49. doi: 10.1186/1754-6834-4-49. PubMed PMID: 22074982; PubMed Central PMCID: PMCPMC3256112.

29. Ding M-Z, Wang X, Yang Y, Yuan Y-J. Comparative metabolic profiling of parental and inhibitors-tolerant yeasts during lignocellulosic ethanol fermentation. Metabolomics. 2012;8(2):232–43. doi: 10.1007/s11306-011-0303-6.

30. Ehrenreich IM, Torabi N, Jia Y, Kent J, Martis S, Shapiro JA, et al. Dissection of genetically complex traits with extremely large pools of yeast segregants. Nature. 2010;464(7291):1039–42. doi: 10.1038/nature08923. PubMed PMID: 20393561; PubMed Central PMCID: PMCPMC2862354.

31. Dimitrov LN, Brem RB, Kruglyak L, Gottschling DE. Polymorphisms in multiple genes contribute to the spontaneous mitochondrial genome instability of Saccharomyces cerevisiae S288C strains. Genetics. 2009;183(1):365–83. doi: 10.1534/genetics.109.104497. PubMed PMID: 19581448; PubMed Central PMCID: PMCPMC2746160.

32. Sardi M, Rovinskiy N, Zhang Y, Gasch AP. Leveraging Genetic-Background Effects in Saccharomyces cerevisiae To Improve Lignocellulosic Hydrolysate Tolerance. Appl Environ Microbiol. 2016;82(19):5838–49. doi: 10.1128/AEM.01603-16. PubMed PMID: 27451446; PubMed Central PMCID: PMCPMC5038035.

33. Cubillos FA, Billi E, Zorgo E, Parts L, Fargier P, Omholt S, et al. Assessing the complex architecture of polygenic traits in diverged yeast populations. Mol Ecol. 2011;20(7):1401–13. doi: 10.1111/j.1365-294X.2011.05005.x. PubMed PMID: 21261765.

34. Carlborg O, Haley CS. Epistasis: too often neglected in complex trait studies? Nat Rev Genet. 2004;5(8):618–25. doi: 10.1038/nrg1407. PubMed PMID: 15266344.

35. Gasch AP, Payseur BA, Pool JE. The Power of Natural Variation for Model Organism Biology. Trends Genet. 2016;32(3):147–54. doi: 10.1016/j.tig.2015.12.003. PubMed PMID: 26777596; PubMed Central PMCID: PMCPMC4769656.

36. Warringer J, Zorgo E, Cubillos FA, Zia A, Gjuvsland A, Simpson JT, et al. Trait variation in yeast is defined by population history. PLoS Genet. 2011;7(6):e1002111. doi: 10.1371/journal.pgen.1002111. PubMed PMID: 21698134; PubMed Central PMCID: PMCPMC3116910.

37. Kvitek DJ, Will JL, Gasch AP. Variations in stress sensitivity and genomic expression in diverse S. cerevisiae isolates. PLoS Genet. 2008;4(10):e1000223. doi: 10.1371/journal.pgen. 1000223. PubMed PMID: 18927628; PubMed Central PMCID: PMCPMC2562515.

38. Wang QM, Liu WQ, Liti G, Wang SA, Bai FY. Surprisingly diverged populations of Saccharomyces cerevisiae in natural environments remote from human activity. Mol Ecol. 2012;21(22):5404–17. doi: 10.1111/j.1365-294X.2012.05732.x. PubMed PMID: 22913817.

39. Bergstrom A, Simpson JT, Salinas F, Barre B, Parts L, Zia A, et al. A high-definition view of functional genetic variation from natural yeast genomes. Mol Biol Evol. 2014;31(4):872–88. doi: 10.1093/molbev/msu037. PubMed PMID: 24425782; PubMed Central PMCID: PMCPMC3969562.

40. Gerke JP, Chen CT, Cohen BA. Natural isolates of Saccharomyces cerevisiae display complex genetic variation in sporulation efficiency. Genetics. 2006;174(2):985–97. doi: 10.1534/genetics.106.058453. PubMed PMID: 16951083; PubMed Central PMCID: PMCPMC1602093.

41. Liti G, Carter DM, Moses AM, Warringer J, Parts L, James SA, et al. Population genomics of domestic and wild yeasts. Nature. 2009;458(7236):337–41. doi: 10.1038/nature07743. PubMed PMID: 19212322; PubMed Central PMCID: PMCPMC2659681.

42. Strope PK, Skelly DA, Kozmin SG, Mahadevan G, Stone EA, Magwene PM, et al. The 100-genomes strains, an S. cerevisiae resource that illuminates its natural phenotypic and genotypic variation and emergence as an opportunistic pathogen. Genome Res. 2015;25(5):762–74. doi: 10.1101/gr.185538.114. PubMed PMID: 25840857; PubMed Central PMCID: PMCPMC4417123.

43. Cromie GA, Hyma KE, Ludlow CL, Garmendia-Torres C, Gilbert TL, May P, et al. Genomic sequence diversity and population structure of Saccharomyces cerevisiae assessed by RAD-seq. G3 (Bethesda). 2013;3(12):2163–71. doi: 10.1534/g3.113.007492. PubMed PMID: 24122055; PubMed Central PMCID: PMCPMC3852379.

44. Zheng YL, Wang SA. Stress Tolerance Variations in Saccharomyces cerevisiae Strains from Diverse Ecological Sources and Geographical Locations. PLoS One. 2015;10(8):e0133889. doi: 10.1371/journal.pone.0133889. PubMed PMID: 26244846; PubMed Central PMCID: PMCPMC4526645.

45. Townsend JP, Cavalieri D, Hartl DL. Population genetic variation in genome-wide gene expression. Mol Biol Evol. 2003;20(6):955–63. doi: 10.1093/molbev/msg106. PubMed PMID: 12716989.

46. Cavalieri D, Townsend JP, Hartl DL. Manifold anomalies in gene expression in a vineyard isolate of Saccharomyces cerevisiae revealed by DNA microarray analysis. Proc Natl Acad Sci U S A. 2000;97(22):12369–74. doi: 10.1073/pnas.210395297. PubMed PMID: 11035792; PubMed Central PMCID: PMCPMC17348.

47. Foss EJ, Radulovic D, Shaffer SA, Ruderfer DM, Bedalov A, Goodlett DR, et al. Genetic basis of proteome variation in yeast. Nat Genet. 2007;39(11):1369–75. doi: 10.1038/ng.2007.22. PubMed PMID: 17952072.

48. Picotti P, Clement-Ziza M, Lam H, Campbell DS, Schmidt A, Deutsch EW, et al. A complete mass-spectrometric map of the yeast proteome applied to quantitative trait analysis. Nature. 2013;494(7436):266–70. doi: 10.1038/nature11835. PubMed PMID: 23334424; PubMed Central PMCID: PMCPMC3951219.

49. Parts L, Liu YC, Tekkedil MM, Steinmetz LM, Caudy AA, Fraser AG, et al. Heritability and genetic basis of protein level variation in an outbred population. Genome Res. 2014;24(8):1363–70. doi: 10.1101/gr.170506.113. PubMed PMID: 24823668; PubMed Central PMCID: PMCPMC4120089.

50. Breunig JS, Hackett SR, Rabinowitz JD, Kruglyak L. Genetic basis of metabolome variation in yeast. PLoS Genet. 2014;10(3):e1004142. doi: 10.1371/journal.pgen. 1004142. PubMed PMID: 24603560; PubMed Central PMCID: PMCPMC3945093.

51. Zhu J, Sova P, Xu Q, Dombek KM, Xu EY, Vu H, et al. Stitching together multiple data dimensions reveals interacting metabolomic and transcriptomic networks that modulate cell regulation. PLoS Biol. 2012;10(4):e1001301. doi: 10.1371/journal.pbio.1001301. PubMed PMID: 22509135; PubMed Central PMCID: PMCPMC3317911 partially funded by Merck.

52. Skelly DA, Merrihew GE, Riffle M, Connelly CF, Kerr EO, Johansson M, et al. Integrative phenomics reveals insight into the structure of phenotypic diversity in budding yeast. Genome Res. 2013;23(9):1496–504. doi: 10.1101/gr.155762.113. PubMed PMID: 23720455; PubMed Central PMCID: PMCPMC3759725.

53. Clowers KJ, Heilberger J, Piotrowski JS, Will JL, Gasch AP. Ecological and Genetic Barriers Differentiate Natural Populations of Saccharomyces cerevisiae. Mol Biol Evol. 2015;32(9):2317–27. doi: 10.1093/molbev/msv112. PubMed PMID: 25953281; PubMed Central PMCID: PMCPMC4540968.

54. Gallone B, Steensels J, Prahl T, Soriaga L, Saels V, Herrera-Malaver B, et al. Domestication and Divergence of Saccharomyces cerevisiae Beer Yeasts. Cell. 2016;166(6):1397–410e16. doi: 10.1016/j.cell.2016.08.020. PubMed PMID: 27610566; PubMed Central PMCID: PMCPMC5018251.

55. Marullo P, Aigle M, Bely M, Masneuf-Pomarede I, Durrens P, Dubourdieu D, et al. Single QTL mapping and nucleotide-level resolution of a physiologic trait in wine Saccharomyces cerevisiae strains. FEMS Yeast Res. 2007;7(6):941–52. doi: 10.1111/j.1567-1364.2007.00252.x. PubMed PMID: 17537182.

56. Meijnen JP, Randazzo P, Foulquie-Moreno MR, van den Brink J, Vandecruys P, Stojiljkovic M, et al. Polygenic analysis and targeted improvement of the complex trait of high acetic acid tolerance in the yeast Saccharomyces cerevisiae. Biotechnol Biofuels. 2016;9:5. doi: 10.1186/s13068-015-0421-x. PubMed PMID: 26740819; PubMed Central PMCID: PMCPMC4702306.

57. Marullo P, Bely M, Masneuf-Pomarede I, Pons M, Aigle M, Dubourdieu D. Breeding strategies for combining fermentative qualities and reducing off-flavor production in a wine yeast model. FEMS Yeast Res. 2006;6(2):268–79. doi: 10.1111/j.1567-1364.2006.00034.x. PubMed PMID: 16487348.

58. Marullo P, Mansour C, Dufour M, Albertin W, Sicard D, Bely M, et al. Genetic improvement of thermo-tolerance in wine Saccharomyces cerevisiae strains by a backcross approach. FEMS Yeast Res. 2009;9(8):1148–60. doi: 10.1111/j.1567-1364.2009.00550.x. PubMed PMID: 19758333.

59. Benjaphokee S, Hasegawa D, Yokota D, Asvarak T, Auesukaree C, Sugiyama M, et al. Highly efficient bioethanol production by a Saccharomyces cerevisiae strain with multiple stress tolerance to high temperature, acid and ethanol. N Biotechnol. 2012;29(3):379–86. doi: 10.1016/j.nbt.2011.07.002. PubMed PMID: 21820088.

60. Hose J, Yong CM, Sardi M, Wang Z, Newton MA, Gasch AP. Dosage compensation can buffer copy-number variation in wild yeast. Elife. 2015;4. doi: 10.7554/eLife.05462. PubMed PMID: 25955966; PubMed Central PMCID: PMCPMC4448642.

61. Cherry JM, Hong EL, Amundsen C, Balakrishnan R, Binkley G, Chan ET, et al. Saccharomyces Genome Database: the genomics resource of budding yeast. Nucleic Acids Res. 2012;40(Database issue):D700–5. doi: 10.1093/nar/gkr1029. PubMed PMID: 22110037; PubMed Central PMCID: PMCPMC3245034.

62. Keane OM, Toft C, Carretero-Paulet L, Jones GW, Fares MA. Preservation of genetic and regulatory robustness in ancient gene duplicates of Saccharomyces cerevisiae. Genome Res. 2014;24(11):1830–41. doi: 10.1101/gr.176792.114. PubMed PMID: 25149527; PubMed Central PMCID: PMCPMC4216924.

63. Dowell RD, Ryan O, Jansen A, Cheung D, Agarwala S, Danford T, et al. Genotype to phenotype: a complex problem. Science. 2010;328(5977):469. doi: 10.1126/science. 1189015. PubMed PMID: 20413493; PubMed Central PMCID: PMCPMC4412269.

64. Schacherer J, Shapiro JA, Ruderfer DM, Kruglyak L. Comprehensive polymorphism survey elucidates population structure of Saccharomyces cerevisiae. Nature. 2009;458(7236):342–5. doi: 10.1038/nature07670. PubMed PMID: 19212320; PubMed Central PMCID: PMCPMC2782482.

65. Palmqvist E, Hahn-Hägerdal B. Fermentation of lignocellulosic hydrolysates. II: inhibitors and mechanisms of inhibition. Bioresource Technology. 2000;74(1):25–33. doi: http://dx.doi.org/10.1016/S0960-8524(99)00161-3·

66. Connelly CF, Akey JM. On the prospects of whole-genome association mapping in Saccharomyces cerevisiae. Genetics. 2012;191(4):1345–53. doi: 10.1534/genetics.112.141168. PubMed PMID: 22673807; PubMed Central PMCID: PMCPMC3416012.

67. Diao L, Chen KC. Local ancestry corrects for population structure in Saccharomyces cerevisiae genome-wide association studies. Genetics. 2012;192(4):1503–11. doi: 10.1534/genetics.112.144790. PubMed PMID: 23023004; PubMed Central PMCID: PMCPMC3512155.

68. Tang Y, Liu X, Wang J, Li M, Wang Q, Tian F, et al. GAPIT Version 2: An Enhanced Integrated Tool for Genomic Association and Prediction. Plant Genome. 2016;9(2). doi: 10.3835/plantgenome2015.11.0120. PubMed PMID: 27898829.

69. Joo JW, Hormozdiari F, Han B, Eskin E. Multiple testing correction in linear mixed models. Genome Biol. 2016;17:62. doi: 10.1186/s13059-016-0903-6. PubMed PMID: 27039378; PubMed Central PMCID: PMCPMC4818520.

70. Ober U, Ayroles JF, Stone EA, Richards S, Zhu D, Gibbs RA, et al. Using whole-genome sequence data to predict quantitative trait phenotypes in Drosophila melanogaster. PLoS Genet. 2012;8(5):e1002685. doi: 10.1371/journal.pgen.1002685. PubMed PMID: 22570636; PubMed Central PMCID: PMCPMC3342952.

71. Westman JO, Mapelli V, Taherzadeh MJ, Franzen CJ. Flocculation causes inhibitor tolerance in Saccharomyces cerevisiae for second-generation bioethanol production. Appl Environ Microbiol. 2014;80(22):6908–18. doi: 10.1128/AEM.01906-14. PubMed PMID: 25172866; PubMed Central PMCID: PMCPMC4249023.

72. Sarvari Horvath I, Franzen CJ, Taherzadeh MJ, Niklasson C, Liden G. Effects of furfural on the respiratory metabolism of Saccharomyces cerevisiae in glucose-limited chemostats. Appl Environ Microbiol. 2003;69(7):4076–86. PubMed PMID: 12839784; PubMed Central PMCID: PMCPMC165176.

73. Guo Z, Olsson L. Physiological response of Saccharomyces cerevisiae to weak acids present in lignocellulosic hydrolysate. FEMS Yeast Res. 2014;14(8):1234–48. doi: 10.1111/1567-1364.12221. PubMed PMID: 25331461.

74. Endo A, Nakamura T, Shima J. Involvement of ergosterol in tolerance to vanillin, a potential inhibitor of bioethanol fermentation, in Saccharomyces cerevisiae. FEMS Microbiol Lett. 2009;299(1):95–9. doi: 10.1111/j.1574-6968.2009.01733.x. PubMed PMID: 19686341.

75. Kodedova M, Sychrova H. Changes in the Sterol Composition of the Plasma Membrane Affect Membrane Potential, Salt Tolerance and the Activity of Multidrug Resistance Pumps in Saccharomyces cerevisiae. PLoS One. 2015;10(9):e0139306. doi: 10.1371/journal.pone.0139306. PubMed PMID: 26418026; PubMed Central PMCID: PMCPMC4587746.

76. Watts T, Khalimonchuk O, Wolf RZ, Turk EM, Mohr G, Winge DR. Mnel is a novel component of the mitochondrial splicing apparatus responsible for processing of a COX1 group I intron in yeast. J Biol Chem. 2011;286(12):10137–46. doi: 10.1074/jbc.M110.205625. PubMed PMID: 21257754; PubMed Central PMCID: PMCPMC3060465.

77. Watari J, Kudo M, Nishikawa N, Kamimura M. Construction of Flocculent Yeast Cells (Saccharomyces cerevisiae) by Mating or Protoplast Fusion Using a Yeast Cell Containing the Flocculation Gene FL05. Agricultural and Biological Chemistry. 1990;54(7):1677–81. doi: 10.1080/00021369.1990.10870222.

78. Jorgensen P, Nelson B, Robinson MD, Chen Y, Andrews B, Tyers M, et al. High-resolution genetic mapping with ordered arrays of Saccharomyces cerevisiae deletion mutants. Genetics. 2002;162(3):1091–9. PubMed PMID: 12454058; PubMed Central PMCID: PMCPMC1462329.

79. Wanat JJ, Singh N, Alani E. The effect of genetic background on the function of Saccharomyces cerevisiae mlh1 alleles that correspond to HNPCC missense mutations. Hum Mol Genet. 2007;16(4):445–52. doi: 10.1093/hmg/dd1479. PubMed PMID: 17210669.

80. Young MJ, Court DA. Effects of the S288c genetic background and common auxotrophic markers on mitochondrial DNA function in Saccharomyces cerevisiae. Yeast (Chichester, England). 2008;25(12):903–12. doi: 10.1002/yea.1644. PubMed PMID: 19160453.

81. Sinha H, Nicholson BP, Steinmetz LM, McCusker JH. Complex genetic interactions in a quantitative trait locus. PLoS Genet. 2006;2(2):e13. doi: 10.1371/journal.pgen.0020013. PubMed PMID: 16462944; PubMed Central PMCID: PMCPMC1359075.

82. Sterken MG, Snoek LB, Kammenga JE, Andersen EC. The laboratory domestication of Caenorhabditis elegans. Trends Genet. 2015;31(5):224–31. doi: 10.1016/j.tig.2015.02.009. PubMed PMID: 25804345; PubMed Central PMCID: PMCPMC4417040.

83. Chandler CH, Chari S, Tack D, Dworkin I. Causes and consequences of genetic background effects illuminated by integrative genomic analysis. Genetics. 2014;196(4):1321–36. doi: 10.1534/genetics.113.159426. PubMed PMID: 24504186; PubMed Central PMCID: PMCPMC3982700.

84. Vu V, Verster AJ, Schertzberg M, Chuluunbaatar T, Spensley M, Pajkic D, et al. Natural Variation in Gene Expression Modulates the Severity of Mutant Phenotypes. Cell. 2015;162(2):391–402. doi: 10.1016/j.cell.2015.06.037. PubMed PMID: 26186192.

85. Glazier AM, Nadeau JH, Aitman TJ. Finding Genes That Underlie Complex Traits. Science. 2002;298(5602):2345–9. doi: 10.1126/science.1076641.

86. Moore JH, Asselbergs FW, Williams SM. Bioinformatics challenges for genome-wide association studies. Bioinformatics. 2010;26(4):445–55. doi: 10.1093/bioinformatics/btp713. PubMed PMID: 20053841; PubMed Central PMCID: PMCPMC2820680.

87. Hu XH, Wang MH, Tan T, Li JR, Yang H, Leach L, et al. Genetic dissection of ethanol tolerance in the budding yeast Saccharomyces cerevisiae. Genetics. 2007;175(3):1479–87. doi: 10.1534/genetics.106.065292. PubMed PMID: 17194785; PubMed Central PMCID: PMCPMC1840089.

88. Parts L, Cubillos FA, Warringer J, Jain K, Salinas F, Bumpstead SJ, et al. Revealing the genetic structure of a trait by sequencing a population under selection. Genome Res. 2011;21(7):1131–8. doi: 10.1101/gr.116731.110. PubMed PMID: 21422276; PubMed Central PMCID: PMCPMC3129255.

89. Katou T, Namise M, Kitagaki H, Akao T, Shimoi H. QTL mapping of sake brewing characteristics of yeast. J Biosci Bioeng. 2009;107(4):383–93. doi: 10.1016/j.jbiosc.2008.12.014. PubMed PMID: 19332297.

90. Wenger JW, Schwartz K, Sherlock G. Bulk segregant analysis by high-throughput sequencing reveals a novel xylose utilization gene from Saccharomyces cerevisiae. PLoS Genet. 2010;6(5):e1000942. doi: 10.1371/journal.pgen.000942. PubMed PMID: 20485559; PubMed Central PMCID: PMCPMC2869308.

91. Liu ZL, Slininger PJ, Dien BS, Berhow MA, Kurtzman CP, Gorsich SW. Adaptive response of yeasts to furfural and 5-hydroxymethylfurfural and new chemical evidence for HMF conversion to 2,5-bis-hydroxymethylfuran. J Ind Microbiol Biotechnol. 2004;31(8):345–52. doi: 10.1007/s10295-004-0148-3. PubMed PMID: 15338422.

92. Nilsson A, Gorwa-Grauslund MF, Hahn-Hagerdal B, Liden G. Cofactor dependence in furan reduction by Saccharomyces cerevisiae in fermentation of acid-hydrolyzed lignocellulose. Appl Environ Microbiol. 2005;71(12):7866–71. doi: 10.1128/AEM.71.12.7866-7871.2005. PubMed PMID: 16332761; PubMed Central PMCID: PMCPMC1317483.

93. Liu ZL, Moon J. A novel NADPH-dependent aldehyde reductase gene from Saccharomyces cerevisiae NRRL Y-12632 involved in the detoxification of aldehyde inhibitors derived from lignocellulosic biomass conversion. Gene. 2009;446(1):1–10. doi: 10.1016/j.gene.2009.06.018. PubMed PMID: 19577617.

94. Almeida JR, Roder A, Modig T, Laadan B, Liden G, Gorwa-Grauslund MF. NADH-vs NADPH-coupled reduction of 5-hydroxymethyl furfural (HMF) and its implications on product distribution in Saccharomyces cerevisiae. Appl Microbiol Biotechnol. 2008;78(6):939–45. doi: 10.1007/s00253-008-1364-y. PubMed PMID: 18330568.

95. Taherzadeh MJ, Gustafsson L, Niklasson C, Liden G. Physiological effects of 5-hydroxymethylfurfural on Saccharomyces cerevisiae. Appl Microbiol Biotechnol. 2000;53(6):701–8. PubMed PMID: 10919330.

96. Bauer BE, Rossington D, Mollapour M, Mamnun Y, Kuchler K, Piper PW. Weak organic acid stress inhibits aromatic amino acid uptake by yeast, causing a strong influence of amino acid auxotrophies on the phenotypes of membrane transporter mutants. Eur J Biochem. 2003;270(15):3189–95. PubMed PMID: 12869194.

97. Hillenmeyer ME, Fung E, Wildenhain J, Pierce SE, Hoon S, Lee W, et al. The chemical genomic portrait of yeast: uncovering a phenotype for all genes. Science. 2008;320(5874):362–5. doi: 10.1126/science.1150021. PubMed PMID: 18420932; PubMed Central PMCID: PMCPMC2794835.

98. Hou L. Novel methods of genome shuffling in Saccharomyces cerevisiae. Biotechnol Lett. 2009;31(5):671–7. doi: 10.1007/s10529-009-9916-5. PubMed PMID: 19153667.

99. Wang H, Hou L. Genome shuffling to improve fermentation properties of top-fermenting yeast by the improvement of stress tolerance. Food Science and Biotechnology. 2010;19(1):145–50. doi: 10.1007/s10068-010-0020-3.

100. Snoek T, Picca Nicolino M, Van den Bremt S, Mertens S, Saels V, Verplaetse A, et al. Large-scale robot-assisted genome shuffling yields industrial Saccharomyces cerevisiae yeasts with increased ethanol tolerance. Biotechnol Biofuels. 2015;8:32. doi: 10.1186/s13068-015-0216-0. PubMed PMID: 25759747; PubMed Central PMCID: PMCPMC4354739.

101. Brachmann CB, Davies A, Cost GJ, Caputo E, Li J, Hieter P, et al. Designer deletion strains derived from Saccharomyces cerevisiae S288C: a useful set of strains and plasmids for PCR-mediated gene disruption and other applications. Yeast (Chichester, England). 1998;14(2):115–32. doi: 10.1002/(SICI)1097-0061(19980130)14:2<115::AID-YEA204>3.0.CO;2-2. PubMed PMID: 9483801.

102. Alexander WG, Doering DT, Hittinger CT. High-efficiency genome editing and allele replacement in prototrophic and wild strains of Saccharomyces. Genetics. 2014;198(3):859–66. doi: 10.1534/genetics.114.170118. PubMed PMID: 25209147; PubMed Central PMCID: PMCPMC4224175.

103. Steinmetz LM, Sinha H, Richards DR, Spiegelman JI, Oefner PJ, McCusker JH, et al. Dissecting the architecture of a quantitative trait locus in yeast. Nature. 2002;416(6878):326–30. doi: 10.1038/416326a. PubMed PMID: 11907579.

104. Xiao W. Yeast protocols: Springer; 2006.

105. Engel SR, Dietrich FS, Fisk DG, Binkley G, Balakrishnan R, Costanzo MC, et al. The reference genome sequence of Saccharomyces cerevisiae: then and now. G3 (Bethesda). 2014;4(3):389–98. doi: 10.1534/g3.113.008995. PubMed PMID: 24374639; PubMed Central PMCID: PMCPMC3962479.

106. Li H. Aligning sequence reads, clone sequences and assembly contigs with BWA-MEM. eprint arXiv:13033997. 2013.

107. McKenna A, Hanna M, Banks E, Sivachenko A, Cibulskis K, Kernytsky A, et al. The Genome Analysis Toolkit: a MapReduce framework for analyzing next-generation DNA sequencing data. Genome Res. 2010;20(9):1297–303. doi: 10.1101/gr. 107524.110. PubMed PMID: 20644199; PubMed Central PMCID: PMCPMC2928508.

108. Danecek P, Auton A, Abecasis G, Albers CA, Banks E, DePristo MA, et al. The variant call format and VCFtools. Bioinformatics. 2011;27(15):2156–8. doi: 10.1093/bioinformatics/btr330. PubMed PMID: 21653522; PubMed Central PMCID: PMCPMC3137218.

109. Cingolani P, Platts A, Wang le L, Coon M, Nguyen T, Wang L, et al. A program for annotating and predicting the effects of single nucleotide polymorphisms, SnpEff: SNPs in the genome of Drosophila melanogaster strain w1118; iso-2; iso-3. Fly (Austin). 2012;6(2):80–92. doi: 10.4161/fly.19695. PubMed PMID: 22728672; PubMed Central PMCID: PMCPMC3679285.

110. Jombart T, Ahmed I. adegenet 1.3-1: new tools for the analysis of genome-wide SNP data. Bioinformatics. 2011;27(21):3070–1. doi: 10.1093/bioinformatics/btr521. PubMed PMID: 21926124; PubMed Central PMCID: PMCPMC3198581.

111. Gao X, Starmer J, Martin ER. A multiple testing correction method for genetic association studies using correlated single nucleotide polymorphisms. Genet Epidemiol. 2008;32(4):361–9. doi: 10.1002/gepi.20310. PubMed PMID: 18271029.

112. Gao X, Becker LC, Becker DM, Starmer JD, Province MA. Avoiding the high Bonferroni penalty in genome-wide association studies. Genet Epidemiol. 2010;34(1):100–5. doi: 10.1002/gepi.20430. PubMed PMID: 19434714; PubMed Central PMCID: PMCPMC2796708.

